# Patient-specific iPSCs carrying an *SFTPC* mutation reveal the intrinsic alveolar epithelial dysfunction at the inception of interstitial lung disease

**DOI:** 10.1101/2020.11.13.382390

**Authors:** Konstantinos-Dionysios Alysandratos, Scott J. Russo, Anton Petcherski, Evan P. Taddeo, Rebeca Acín-Pérez, Carlos Villacorta-Martin, J. C. Jean, Surafel Mulugeta, Benjamin C. Blum, Ryan M. Hekman, Marall Vedaie, Seunghyi Kook, Jennifer A. Wambach, F. Sessions Cole, Aaron Hamvas, Andrew Emili, Susan H. Guttentag, Orian S. Shirihai, Michael F. Beers, Darrell N. Kotton

**Affiliations:** Center for Regenerative Medicine, Boston University and Boston Medical Center, Boston, MA 02118, USA; The Pulmonary Center and Department of Medicine, Boston University School of Medicine, Boston, MA 02118, USA; Pulmonary, Allergy, and Critical Care Division, Department of Medicine, University of Pennsylvania Perelman School of Medicine, Philadelphia, PA 19104, USA; PENN-CHOP Lung Biology Institute, University of Pennsylvania Perelman School of Medicine, Philadelphia, PA 19104, USA; Departments of Medicine, Endocrinology and Molecular and Medical Pharmacology, David Geffen School of Medicine at UCLA, Los Angeles, CA 90095, USA; Departments of Biology and Biochemistry, Boston University School of Medicine, Boston, MA 02118, USA; Department of Pediatrics; Monroe Carell Jr. Children’s Hospital, Vanderbilt University, Nashville, TN 37232, USA; Division of Newborn Medicine, Edward Mallinckrodt Department of Pediatrics, Washington University School of Medicine and St. Louis Children’s Hospital, St. Louis, MO 63110, USA; Department of Pediatrics, Northwestern University Feinberg School of Medicine, Chicago, IL 60611, USA

## Abstract

The incompletely understood pathogenesis of pulmonary fibrosis (PF) and lack of reliable preclinical disease models have limited development of effective therapies. An emerging literature now implicates alveolar epithelial type 2 cell (AEC2) dysfunction as an initiating pathogenic event in the onset of a variety of PF syndromes, including adult idiopathic pulmonary fibrosis (IPF) and childhood interstitial lung disease (chILD). However, inability to access primary AEC2s from patients, particularly at early disease stages, has impeded identification of disease-initiating mechanisms. Here we present an *in vitro* reductionist model system that permits investigation of epithelial-intrinsic events that lead to AEC2 dysfunction over time using patient-derived cells that carry a disease-associated variant, *SFTPC^I73T^*, known to be expressed solely in AEC2s. After generating patient-specific induced pluripotent stem cells (iPSCs) and engineering their gene-edited (corrected) counterparts, we employ directed differentiation to produce pure populations of syngeneic corrected and mutant AEC2s, which we expand >10^15^ fold *in vitro*, providing a renewable source of cells for modeling disease onset. We find that mutant iPSC-derived AEC2s (iAEC2s) accumulate large amounts of misprocessed pro-SFTPC protein which mistrafficks to the plasma membrane, similar to changes observed *in vivo* in the donor patient’s AEC2s. These changes result in marked reduction in AEC2 progenitor capacity and several downstream perturbations in AEC2 proteostatic and bioenergetic programs, including a late block in autophagic flux, accumulation of dysfunctional mitochondria with consequent time-dependent metabolic reprograming from oxidative phosphorylation to glycolysis, and activation of an NF-κB dependent inflammatory response. Treatment of *SFTPC^I73T^* expressing iAEC2s with hydroxychloroquine, a medication commonly prescribed to these patients, results in aggravation of autophagy perturbations and metabolic reprogramming. Thus, iAEC2s provide a patientspecific preclinical platform for modeling the intrinsic epithelial dysfunction associated with the inception of interstitial lung disease.

## Introduction

Idiopathic pulmonary fibrosis (IPF) is the most common and severe form of idiopathic interstitial pneumonia and is characterized by relentless fibrosis leading to disruption of the gas exchange unit and death within an average of four years from the time of diagnosis (Lederer and Martinez, 2018; Raghu et al., 2016; 2014; Travis et al., 2013). The poorly understood pathogenesis of IPF, in part due to the lack of human models of the disease, has been a major hurdle in developing effective therapies, and the two currently FDA-approved drugs pirfenidone and nintedanib, both aimed at inhibiting fibrogenic fibroblasts, have been shown to slow but not reverse the progression of lung function decline (King et al., 2014; Richeldi et al., 2014).

While a broad, established literature has focused on the role of lung fibroblasts in perpetuating IPF at later disease stages, factors that initiate or drive disease onset, particularly at early stages, have been particularly hard to identify. With the advent of genome wide association studies and intensive study of familial forms of pulmonary fibrosis, the lung epithelium has been increasingly implicated as a potential proximal disease driver, with variants in gene loci expressed in lung epithelia having been associated with disease risk (C. K. Garcia, 2018; Kropski et al., 2015).

Of the many types of epithelia present in the lung, dysfunction of the alveolar epithelial type 2 cell (AEC2), in particular, has been repeatedly implicated in the pathogenesis of a variety of interstitial lung disease (ILD) syndromes, including IPF (Barkauskas and Noble, 2014; Katzenstein, 1985; Nichelle I Winters MD et al., 2019; Selman and Pardo, 2014), with the most compelling evidence arising from genetic studies revealing an association between mutations in AEC2-specific genes and the development of pulmonary fibrosis (C. K. Garcia, 2018; Kropski et al., 2015). Determining how AEC2 dysfunction leads to disease in humans, however, has been challenging to date. For example, studying AEC2 pathophysiology in primary human cells has been limited by difficulty in accessing patient samples and the tendency of AEC2s to lose their identity in culture (Borok et al., 1998; Foster et al., 2007), in the absence of mesenchymal feeders (Barkauskas et al., 2013).

An ideal model of human AEC2 dysfunction would allow the study of disease by utilizing patient-derived cells to reveal a cascade of mechanistic events associated with the inception as well as progression of lung disease. Here, we present such a model of disease inception employing induced pluripotent stem cells (iPSCs) we have derived from a patient with ILD in order to generate an inexhaustible supply of AEC2-like cells *in vitro* to study the epithelial-intrinsic events that lead to specific AEC2 perturbations that are then validated in the donor patient’s tissue *in vivo*. To ensure examination of only the cell type responsible for initiating disease without the potential confounding or secondary effects of co-cultured supporting cells, we have developed an epithelial-only model composed solely of purified AEC2s that can be propagated indefinitely without any supporting cells, and we have chosen to focus here on a disease-associated variant in the surfactant protein C *(SFTPC)* gene, since it is known to be expressed postnatally solely in AEC2s and can be gene edited to provide syngeneic comparator AEC2s, thus providing a reductionist model amenable to discerning the putative epithelial-intrinsic events associated with disease inception.

Heterozygous mutations in *SFTPC* have been associated with both sporadic and familial pulmonary fibrosis as well as childhood interstitial lung disease (chILD) (Cottin et al., 2011; Crossno et al., 2010; Mulugeta et al., 2015; Nogee et al., 2001; Ono et al., 2011; Thomas et al., 2002; van Moorsel et al., 2010). Among the more than 60 *SFTPC* variants described to date, the missense mutation g.1286T>C resulting in substitution of threonine for isoleucine at amino acid 73 in the pro-SFTPC protein (SFTPC^I73T^), is the focus of our study because it is the predominant disease-associated *SFTPC* variant. Two recent mouse genetic models, conditionally expressing mutant SFTPC (p.I73T and p.C121G, respectively) from the endogenous *SFTPC* locus, resulted in polycellular alveolitis and pulmonary fibrosis (Katzen et al., 2019; Nureki et al., 2018). While previous studies have shown that expression of *SFTPC^I73T^ in vitro* (Beers et al., 2011; A. Hawkins et al., 2015) or in mice *in vivo* (Nureki et al., 2018) leads to SFTPC proprotein misprocessing as well as lung fibrosis, the cellular mechanisms by which dysfunctional AEC2s initiate the fibrotic cascade in humans remain elusive. Most importantly, elucidating the molecular pathogenesis of human AEC2 dysfunction caused by the *SFTPC^I73T^* mutation is likely to inform the broader mechanisms by which AEC2 dysfunction leads to IPF.

A critical role for AEC2s in initiating a variety of interstitial lung diseases is concordant with their essential role in maintaining distal lung homeostasis. AEC2s are highly specialized cells that function both as facultative progenitors of alveoli as well as secretory cells in the distal lung. In order to meet the high metabolic demands posed by these functions, AEC2s have a larger number of mitochondria compared to other lung cell types (Massaro et al., 1975), and alterations in mitochondrial function, as observed in our model, are likely to lead to a variety of perturbed metabolic programs and stress of this key lung cell. Indeed, accumulation of dysmorphic and dysfunctional mitochondria in association with impaired autophagy and mitophagy has been shown in AEC2s from IPF lungs (Bueno et al., 2015). Furthermore, thyroid hormone-mediated restoration of mitochondrial function and mitophagy was recently shown to blunt pulmonary fibrosis in two mouse models of pulmonary fibrosis (Yu et al., 2018).

Since there are significant differences between murine and human pulmonary fibrosis, there is a pressing need for reliable human preclinical disease models that would not only provide further insight into the pathophysiology of human AEC2 dysfunction at the inception of pulmonary fibrosis, but would also serve as a platform to test the safety and efficacy of unproven currently used as well as novel therapies. We here show that *SFTPC^I73T^*-expressing human iPSC-derived AEC2s (hereafter iAEC2s) provide further insights into disease pathogenesis, allow assessment of currently used unproven treatments, and have the potential to serve as a platform for testing novel therapeutics.

## Results

### Generation of patient-specific iPSC lines and their differentiation to alveolar epithelium

To examine the mechanisms leading to human AEC2 dysfunction, we generated a novel patient-specific iPSC line by reprogramming fibroblasts from a patient heterozygous for the most frequent disease-associated *SFTPC* variant (*SFTPC^I73T/WT^*) (Figure 1A). The patient suffered from severe interstitial lung disease (Figure 1B) necessitating the need for lung transplantation in early adolescence (Figure 1C). To overcome the hurdles imposed by the high variance in differentiation efficiencies or cellular phenotypes associated with different genetic backgrounds of iPSC lines generated from distinct individuals (Bock et al., 2011; Boulting et al., 2011; Kim et al., 2010) and to allow for purification of specific cell types, we engineered gene-corrected syngeneic lines by inserting a tdTomato fluorescent reporter into one allele of the endogenous *SFTPC* locus of the parental iPSC line. We used transcription activator-like effector nucleases (TALEN)-based gene editing to knock-in the tdTomato reporter to the endogenous *SFTPC* locus at the start codon, resulting in the generation of syngeneic corrected (*SFTPC^tdTomato/WT^*) and mutant (*SFTPC^I73T/tdTomato^*) iPSC lines, as the tdTomato cassette is followed by a stop/polyA cassette, preventing expression of the subsequent *SFTPC* coding sequence from the targeted allele (Figure 1D).Thus, this targeting strategy was designed to result in the preparation of a reductionist model where only one *SFTPC* allele would be expressed in each paired clone, enabling study of only the mutant vs. only the normal allele. This targeting of the tdTomato reporter into the *SFTPC* locus would enable not only the tracking of SFTPC^tdTomato+^ cells as they emerge through *in vitro* directed differentiation but also the sorting of a pure population of corrected and mutant SFTPC^tdTomato+^ putative iAEC2s to be profiled transcriptomically and functionally, head-to-head.

**Figure 1.**
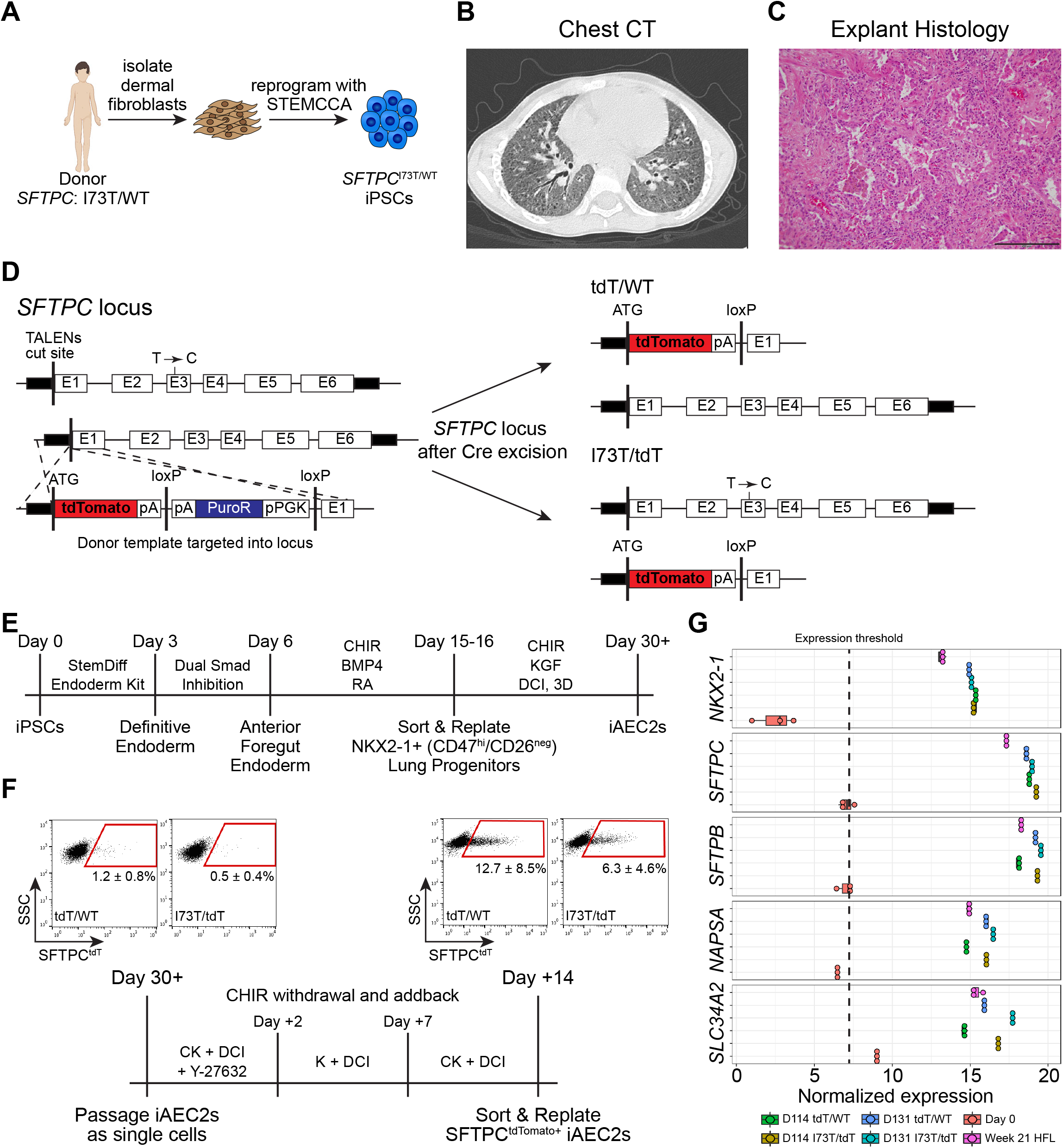
Generation of patient-specific iPSC lines and their differentiation to alveolar epithelium. (A) Schematic showing the generation of patient-specific iPSCs from dermal fibroblasts from a patient carrying the most frequent *SFTPC* pathogenic variant (*SFTPC^I73T/WT^*). (B) Chest CT of the same patient reveals bilateral diffuse ground glass opacities and traction bronchiectasis. (C) H&E staining of the explanted lungs shows end-stage lung disease with interstitial fibrosis, chronic inflammation, and alveolar remodeling as well as AEC2 hyperplasia and degenerating macrophages within the residual alveoli. Scale bar: 200 μm. (D) Transcription activator-like effector nucleases (TALEN) targeting strategy and edited *SFTPC* loci post Cre-mediated antibiotic cassette excision. (E) Schematic of directed differentiation protocol from iPSCs to day 30+ monolayered epithelial iAEC2 spheres (“alveolospheres”). (F) Schematic of CHIR withdrawal and addback to achieve iAEC2 maturation and representative flow cytometry dot plots (mean ± SD is shown; n=6 biological replicates of independent differentiations). (G) Dot plots depicting the normalized expression level of AEC2-marker genes in day 114 and day 131 SFTPC^tdT/WT^ and SFTPC^I73T/tdT^ iAEC2s compared to day 0 iPSCs and week 21 human fetal distal lung (HFL) controls, by bulk RNA sequencing (boxplots represent mean ± SD; n=3 biological replicates of independent wells of a differentiation).

To ensure that any findings from this single allele expression model could be replicated in iAEC2s that express two active *SFTPC* alleles, as normally occurs *in vivo*, we also engineered an untargeted clone of the same patient line (*SFTPC*^I73T/WT^), a version that then underwent footprint-free correction of the *SFTPC* mutation via CRISPR/Cas9 gene editing (*SFTPC*^WT/WT^) (Supplemental Figure 1A).

### Derivation of parallel indefinitely self-renewing iAEC2s

Next, we used our previously published lung directed differentiation protocol (Jacob et al., 2017; 2019) to generate corrected and mutant iAEC2s from iPSCs (Figure 1E), with the goal of producing indefinitely self-renewing patient-specific cells for disease modeling. In order to increase SFTPC expression frequencies, prior to SFTPC^tdTomato+^ cell sorting, iAEC2s underwent a step of transient Wnt signaling modulation (via CHIR withdrawal followed by CHIR addback), which resulted in augmentation of SFTPC^tdTomato^ expression, as expected based on our prior work demonstrating that Wnt downregulation promotes iAEC2 maturation (Jacob et al., 2019) (Figure 1F). Corrected and mutant SFTPC^tdTomato+^ iAEC2s were then sorted to purity for further replating, characterization of gene expression (Figure 1G), and culture expansion as self-renewing epithelial-only spheres without further cell sorting (Figure 2A, B). The resulting iAEC2s maintained an AEC2 specific transcriptomic profile (Figure 1G); however, alveolospheres generated from mutant iAEC2s demonstrated distinct morphologies and brighter tdTomato reporter expression (Figure 2A, B and further discussed below).

**Figure 2.**
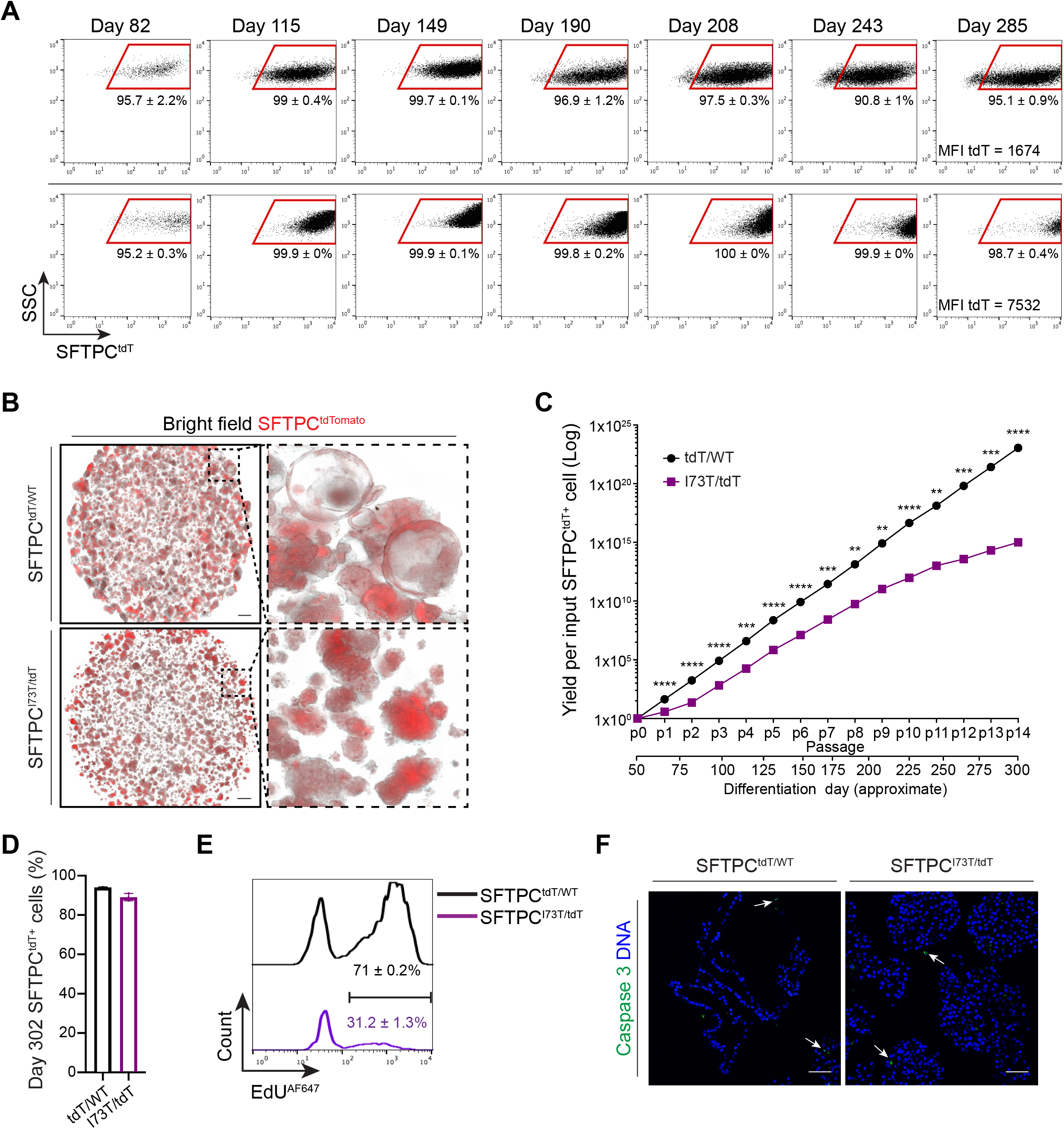
Derivation of parallel self-renewing corrected (SFTPC^tdT/WT^) and mutant (SFTPC^I73T/tdT^) iAEC2s. (A) Representative flow cytometry dot plots of day 82 (p2), day 115 (p4), day 149 (p6), day 190 (p8), day 208 (p9), day 243 (p11), and day 285 (p13) corrected (tdT/WT) and mutant (I73T/tdT) iAEC2s (mean ± SD is shown; n=3 biological replicates of independent wells of a differentiation). MFI: mean fluorescence intensity. (B) Representative live-cell imaging of SFTPC^tdT/WT^ and SFTPC^I73T/tdT^ alveolospheres (bright-field/tdTomato overlay; day 149). Scale bars: 500 μm. (C) Graph showing yield in cell number per input corrected (tdT/WT) or mutant (I73T/tdT) SFTPC^tdTomato+^ sorted cell on day 51. SFTPC^tdTomato+^ yield at each passage is shown without further sorting (mean ± SD; n=3 biological replicates of independent wells of a differentiation). **p<0.01, ***p<0.001, ****p<0.0001 by unpaired, two-tailed Student’s t-test. (D) Bar graph shows retention of the AEC2 cell fate in SFTPC^tdT/WT^ and SFTPC^I73T/tdT^ iAEC2s maintained in culture for 302 days, measured by flow cytometry as the frequency of cells expressing the SFTPC^tdTomato^ reporter (mean ± SD; n=3 biological replicates of independent wells of a differentiation). (E) Histograms show higher proliferation rates in SFTPC^tdT/WT^ compared to SFTPC^I73T/tdT^ iAEC2s, measured by flow cytometry as the frequency of SFTPC^tdTomato+^ cells that incorporate EdU (mean ± SD; n=3 biological replicates of independent wells of a differentiation). (F) Representative confocal immunofluorescence microscopy of SFTPC^tdT/WT^ and SFTPC^I73T/tdT^ iAEC2s stained for activated caspase 3 (green) and DNA (Hoechst, blue) shows absence of significant apoptosis. Scale bars: 50 μm.

We sought to determine both the stability of SFTPC expression levels as well as the total cell yield arising during serial passaging of mutant vs. corrected iAEC2s in epithelial-only 3D cultures. Over a 302 day period we generated yields of >10^23^ and >10^15^ corrected vs. mutant iAEC2s per starting sorted tdTomato+ cell, respectively (Figure 2C). Although yields over this time period were significantly lower for mutant iAEC2s, both lines maintained SFTPC^tdTomato^ expression in 94 ± 0.3% and 89.1 ± 1.9% of cells for at least 302 days and maintained a normal karyotype until at least day 214 (Figure 2D). In this self-renewing model, the mutant iAEC2s demonstrated significantly lower proliferation rates than corrected iAEC2s when assessed by EdU incorporation assays (Figure 2E) and no differences in apoptosis, as measured by immunostaining for activated caspase-3 (Figure 2F). These results suggest SFTPC mutant iAEC2s have diminished selfrenewal capacity.

### SFTPC^I73T^ iAEC2s display aberrant protein trafficking and accumulate large amounts of aberrantly processed SFTPC pro-protein

We further examined the impact of the *SFTPC^I73T^* mutation on AEC2 morphology *in vitro* and *in vivo*. While the corrected (*SFTPC^tdTomato/WT^*) iAEC2s formed monolayered epithelial spheres (alveolospheres) in culture as expected (Jacob et al., 2017), mutant (*SFTPC^I73T/tdTomato^*) iAEC2s lacked lumens and instead formed ball-like structures when assessed by microscopy of paraffin tissue sections (Figure 3A, panel i), semi-thin plastic sections (Figure 3A, panel ii), or EPCAM immunostained sections (Figure 3B). Similarly, we found morphologic changes in developing AEC2s *in vivo* in *SFTPC*^I73T^ knock-in mice, which do not survive birth (Nureki et al., 2018). For example, we examined E18.5 lungs from these mice and found they displayed arrested lung morphogenesis in late sacculation and complete obliteration of primordial saccules by clumps of HA-tagged pro-SFTPC^I73T^ expressing cells (Figure 3C). Analysis of the explanted distal lung from the patient from whom the iPSCs had been derived also revealed distal lung morphological changes with areas of hyperplastic or clumped AEC2s (Figure 3D; arrowheads) that differed from normal lung control tissue sections (Figure 3D; arrowheads). To screen for potential morphological differences in lamellar bodies between mutant and corrected iAEC2s, we employed toluidine blue stained plastic sections (Figure 3A, panel ii) as well as transmission electron microscopy (TEM) (Figure 3E). We observed small inclusions reminiscent of lamellar bodies, present intracellularly in iAEC2s generated from both iPSC lines (Figure 3A panel ii; arrowheads) and extracellularly in the lumens of corrected alveolospheres, possibly reflecting lamellar body exocytosis (Figure 3A panel ii; ellipse), in keeping with our prior publication characterizing surfactant secretion from iAEC2s (Jacob et al., 2017). TEM confirmed these inclusions to be lamellar bodies (Figure 3E), with no appreciable difference in lamellar body ultrastructural morphology observed between mutant and corrected cells, despite the clumped growth patterns observed in mutant cells.

**Figure 3.**
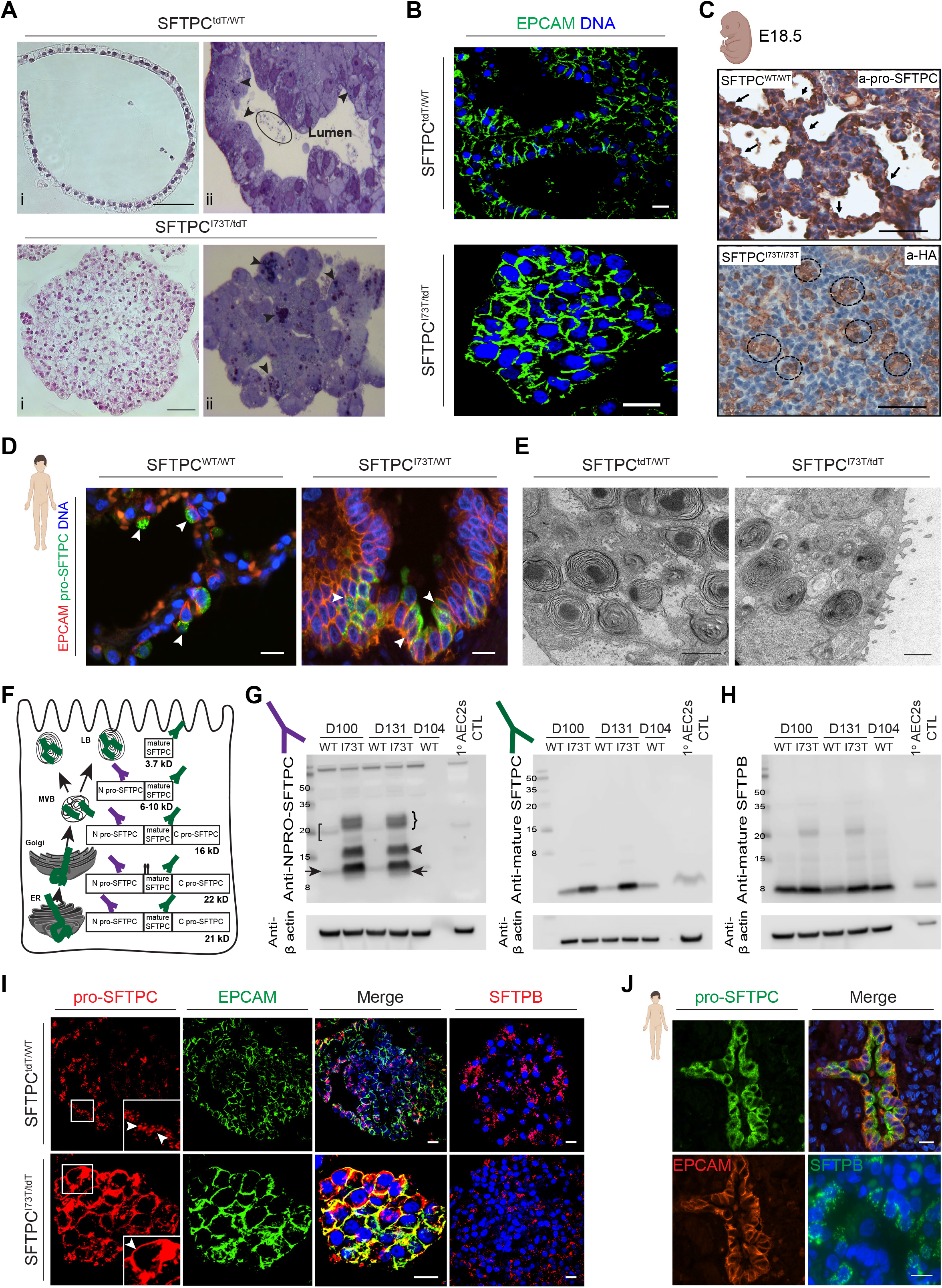
SFTPC^I73T/tdT^ iAEC2s demonstrate distinct cellular morphology and misprocess and mistraffick pro-SFTPC similarly to in vivo SFTPC^I73T^ expressing iAEC2s. (A) Representative H&E staining of formalin fixed and paraffin embedded sections (i, scale bars: 50 μm) and toluidine blue staining of plastic sections (ii) of SFTPC^tdT/WT^ and SFTPC^I73T/tdT^ iAEC2 cultures reveal distinct morphologies. Arrowheads indicate putative lamellar body-like inclusions, circle indicates intraluminal inclusions. (B) Representative confocal immunofluorescence microscopy of SFTPC^tdT/WT^ and SFTPC^I73T/tdT^ iAEC2 cultures for EPCAM (green) and DNA (Hoechst, blue). Scale bars: 10 μm. (C) Representative staining of E18.5 CFlp-SFTPC^I73T/I73T^ embryos for HA-tagged SFTPC reveals tufts of HA+ AEC2s within obliterated saccules compared to CFlp-SFTPC^WT/WT^ controls that show normal developing saccules lined by pro-SFTPC^+^ cells. Scale bars: 70 μm. (D) Representative confocal immunofluorescence microscopy of distal sections of the explanted patient’s lung and healthy distal lung sections stained for EPCAM (red), pro-SFTPC (green), and DNA (Hoechst, blue). Scale bars: 10 μm. (E) Representative TEM images of SFTPC^tdT/WT^ and SFTPC^I73T/tdT^ iAEC2s reveal the presence of lamellar bodies. Scale bars: 1 μm. (F) Schematic depicting the cellular compartments in which pro-SFTPC processing into mature SFTPC occurs. ER: endoplasmic reticulum, MVB: multivesicular body, LB: lamellar body. (G-H) Representative Western blots of SFTPC^tdT/WT^ and SFTPC^I73T/tdT^ iAEC2 lysates at the indicated time points were compared to freshly isolated primary human AEC2s lysates for pro-SFTPC (NPRO-SFTPC) (G, left panel), mature SFTPC (G, right panel), and mature SFTPB (H) with β actin as a loading control. SFTPC^I73T/tdT^ iAEC2s accumulate large amounts of the primary translation product (right bracket) and misprocessed intermediate forms (arrow head). In SFTPC^tdT/WT^ iAEC2s, both the primary translation product (left bracket) and major processed intermediate forms (arrow) are detected. SFTPC^I73T/tdT^ iAEC2s accumulate a larger amount of both mature 3.7 kDa SFTPC and mature 8 kDa SFTPB. (I) Representative confocal immunofluorescence microscopy of SFTPC^tdT/WT^ and SFTPC^I73T/tdT^ iAEC2 cultures for pro-SFTPC (red), EPCAM (green), SFTPB (red), and DNA (Hoechst, blue) reveals pro-SFTPC expression on SFTPC^I73T/tdT^ iAEC2 plasma membranes (inset, arrowhead). In SFTPC^tdT/WT^ iAEC2s, pro-SFTPC is expressed in cytosolic vesicles (inset, arrowheads). SFTPB is expressed in cytosolic vesicles of both SFTPC^tdT/WT^ and SFTPC^I73T/tdT^ iAEC2s. Scale bars: 10 μm. (J) Representative confocal immunofluorescence microscopy of distal sections of the explanted patient lung for pro-SFTPC (green), EPCAM (red), SFTPB (green), and DNA (Hoechst, blue) shows pro-SFTPC expression on AEC2 plasma membranes while SFTPB is expressed in cytosolic vesicles. Scale bars: 10 μm.

To examine the effects of the *SFTPC^I73T^* mutation on human surfactant protein processing, next we analyzed protein extracts from normal primary adult AEC2s in comparison to corrected vs. mutant iAEC2s. In humans SFTPC protein is produced exclusively by AEC2s as a 21 kDa proprotein which undergoes four endoproteolytic cleavages to generate the mature 3.7 kDa form for secretion (Figure 3F) (Beers et al., 1994; Beers and Mulugeta, 2005; Brasch et al., 2002). Complete biosynthesis requires directed anterograde trafficking from the Golgi to lamellar bodies and only cells expressing these organelles are capable of fully processing SFTPC to its mature form. Western blot analysis revealed that the mutant iAEC2s accumulated both significant amounts of misprocessed pro-SFTPC forms (Figure 3G; arrowhead) as well as increased amounts of incompletely processed pro-SFTPC intermediate forms (Figure 3G; right arrow) suggesting aberrant post-Golgi targeting coupled with inefficient posttranslational processing of the SFTPC^I73T^ primary translation product and intermediates (Figure 3G; right bracket and arrow respectively). In contrast, corrected iAEC2s demonstrated restoration of a normal pro-SFTPC processing pattern (Figure 3G: left bracket and arrow) and production of mature 3.7 kDa SFTPC (Figure 3G, right panel). Despite inefficient turnover of mutant pro-SFTPC^I73T^, production of the mature 3.7 kDa SFTPC was also observed in mutant iAEC2s but accumulated to a greater degree than in corrected iAEC2s suggesting its turnover is also inefficient (Figure 3G, right panel). By comparison, the mutant iAEC2s demonstrated intact SFTPB protein processing generating roughly similar amounts of mature 8 kDa SFTPB (Figure 3H), suggesting that differences in mutant SFTPC posttranslational processing and turnover were likely not a result of a broader AEC2 trafficking dysfunction. In support of this, immunostaining revealed normal pro-SFTPC intracellular localization to lamellar bodies of corrected iAEC2s, observed as distinct cytoplasmic puncta, while in the mutant iAEC2s pro-SFTPC was mislocalized to the plasma membrane (Figure 3I). Mistrafficking of pro-SFTPC to the plasma membrane was also demonstrated *in vivo* in sections of the explanted lung of the patient from whom the iPSC line was derived (Figure 3D and J). Similar to the normal SFTPB processing profile, immunostaining revealed normal intracellular localization of SFTPB in both mutant and corrected iAEC2s *in vitro* (Figure 3I) as well as in the donor patient’s lung tissue *in vivo* (Figure 3J). Misprocessing of pro-SFTPC was also validated in the patient line biallelically expressing both the mutant and “footprint-free” corrected *SFTPC* alleles (*SFTPC*^I73T/WT^) (Supplemental Figure 1B).

### Transcriptomic and proteomic profiling of SFTPC^I73T^ iAEC2s and their gene-edited corrected counterparts identifies differential regulation of the lysosomal/autophagy pathway

We next sought to identify the downstream consequences of pro-SFTPC misprocessing and mistrafficking by profiling the mutant and corrected iAEC2s by single cell RNA sequencing (scRNA-seq), bulk RNA sequencing, and proteomic analyses. Transcriptomes profiled at single cell resolution were more similar at early time points (day 30) after the initial onset of *SFTPC* expression but diverged over time at later time points (day 113 or day 114) in two repeated experiments (Figure 4A and Supplemental Figure 3A). For example, only 224 genes were differentially expressed at day 30, whereas 1107 genes were differentially expressed by day 113 between mutant and corrected iAEC2s (FDR <0.05). Cell populations at each time point remained similarly pure in terms of the frequencies of expression of *NKX2-1, SFTPB* (Figure 4B and Supplemental Figure 3B), *SFTPC* (Figure 4C, D, and Supplemental Figure 3B, C), as well as the AEC2 program (Figure 4D and Supplemental Figure 3B), based on quantification of expression of our previously defined 8-gene AEC2 differentiation benchmark established using primary adult AEC2 cells (Hurley et al., 2020). Importantly, the cells did not detectably assume alternative lung or non-lung fates based on little to no expression of airway (*SCGB1A1, FOXJ1, KRT5*; Figure 4B), endothelial (*CDH5*), hematopoietic (*PTPRC*), or non-lung endodermal (*CDX2, AFP, ALB*; Figure 4B, and supplemental figure 3C) transcripts.

**Figure 4.**
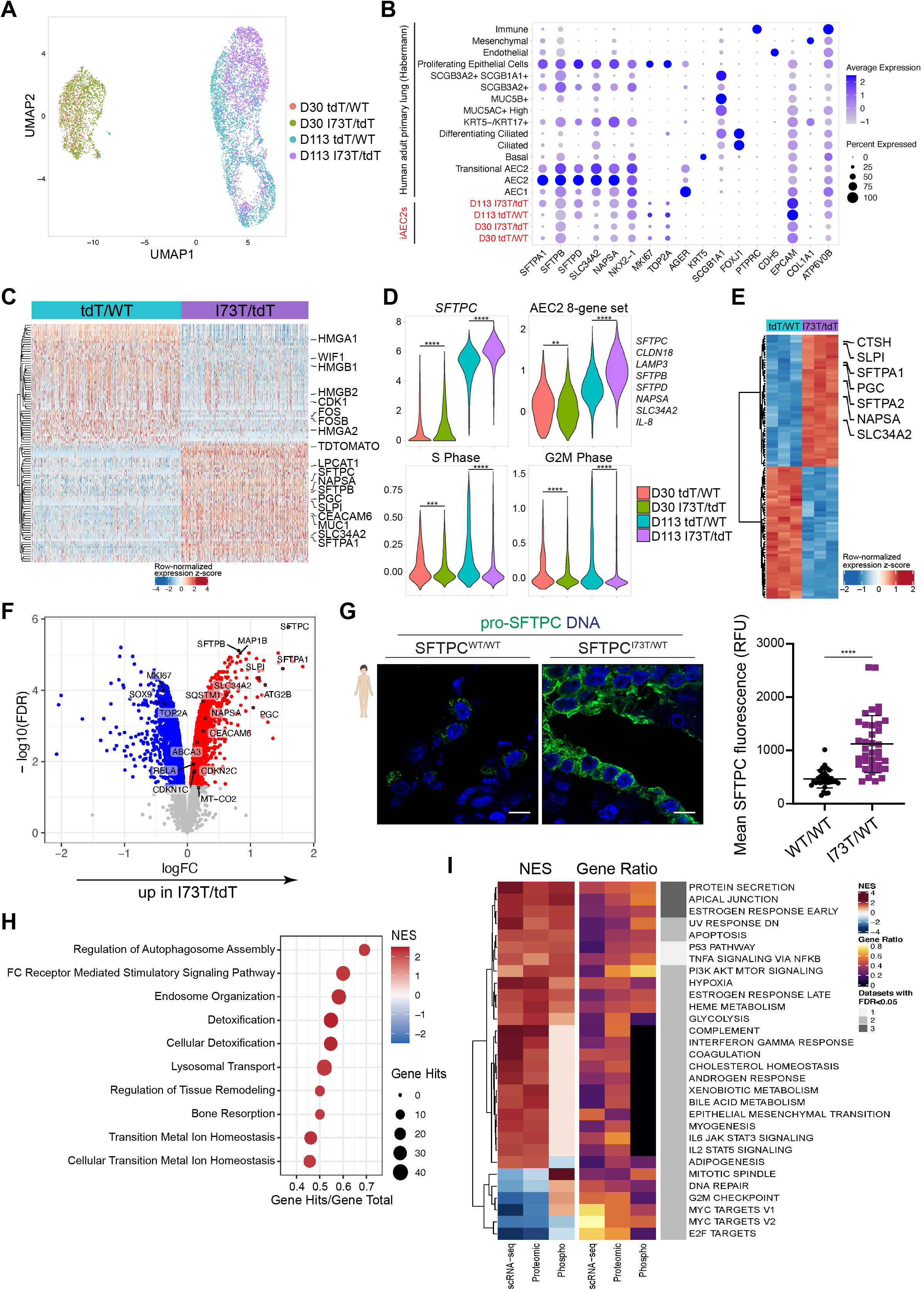
Transcriptomic and proteomic/phosphoproteomic analyses identify candidate disease-associated pathways. (A) Clustering of scRNA-seq transcriptomes at two time points (day 30 and day 113 of differentiation) using Uniform Manifold Approximation Projection (UMAP). Cells are colored based on iAEC2 mutant vs. corrected identity as indicated. (B) Average expression levels and frequencies (purple dots) for selected genes profiled by scRNA-seq in SFTPC^tdT/WT^ and SFTPC^I73T/tdT^ iAEC2s. Comparison is made to a publicly available adult primary distal lung dataset (Habermann et al., 2020) and genes are selected to indicate AEC2, AEC1, airway, endothelial, epithelial, leukocyte, or proliferation programs. (C) Heatmap of the top 50 genes upregulated and the top 50 downregulated comparing day 113 SFTPC^tdT/WT^ vs day 113 SFTPC^I73T/tdT^ iAEC2s by scRNA-seq (ranked by average log fold-change, FDR <0.05; row-normalized expression z-scores are indicated). AEC2-marker genes are highlighted with large font. (D) Violin plots showing normalized expression for indicated genes or cell cycle phase in day 30 and day 113 SFTPC^tdT/WT^ and SFTPC^I73T/tdT^ iAEC2s by scRNA-seq. (E) Heatmap of the top 50 genes upregulated and the top 50 downregulated in day 114 SFTPC^tdT/WT^ vs. day 114 SFTPC^I73T/tdT^ iAEC2s by bulk RNA seq (ranked by FDR, FDR <0.05; row-normalized expression z-scores are indicated). AEC2-marker genes are highlighted with large font. (F) Volcano plots indicating differential protein expression in day 113 SFTPC^tdT/WT^ vs. day 113 SFTPC^I73T/tdT^ iAEC2s. (G) Representative confocal immunofluorescence microscopy of distal sections of the explanted patient’s lung and healthy distal lung sections stained for pro-SFTPC (green) and DNA (Hoechst, blue) shows an altered cellular localization pattern and a higher pro-SFTPC fluorescence intensity in the patient’s AEC2s, quantified by relative fluorescence units (RFU). Scale bars: 10 μm. (mean ± SD; n=35 individual SFTPC-positive cells treated as biological replicates). (H) Top 10 upregulated pathways in day 113 SFTPC^I73T/tdT^ vs. day 113 SFTPC^tdT/WT^ iAEC2s based on global proteomic analysis. (I) Integrated gene set enrichment analyses based on scRNA-seq transcriptomic, proteomic, and phosphoproteomic analyses in day 113 SFTPC^I73T/tdT^ vs. day 113 SFTPC^tdT/WT^ iAEC2s with an FDR<0.1 in at least 2 out of 3 datasets. NES: normalized enrichment score; gene ratio: ratio of enriched genes for a given pathway to the total number of genes in the pathway. The last column represents the number of datasets (out of 3) with an FDR<0.05. **p<0.01, ***p<0.001, ****p<0.0001 by unpaired, two-tailed Student’s t-test for all panels.

Although both mutant and corrected iAEC2s captured for scRNA-Seq analysis expressed similar *frequencies* of the AEC2 program, mutant cells exhibited significantly higher expression *levels* of AEC2 marker genes (Figure 4B-E) and less proliferation (significantly lower MKI67 expression and lower frequencies of cells in G2/S/M phases of cell cycle; Figure 4B, D and Supplemental Figure 3B). For example, 11 out of the top 50 differentially upregulated genes in the mutant iAEC2s in the day 113 scRNA-seq were composed of transcripts encoding surfactants, lamellar body-related, and other AEC2-marker genes (*PGC, SFTPB, SLPI, NAPSA, SLC34A2, LPCAT1, SFTPA1, CEACAM6, MUC1, SFTPC, TDTOMATO*) (Figure 4C). We validated these findings by qRT-PCR quantitation (Supplemental Figure 3D) as well as in independent experiments analyzed by bulk RNA sequencing at 2 time points, day 114 and day 131 of differentiation, again finding AEC2 gene markers comprised 14% of the top 50 most upregulated genes in mutant iAEC2s (Figure 4E).

By global proteomic analysis performed on the same day as scRNA-seq, SFTPC was the most enriched protein overall in mutant vs. corrected iAEC2s (ranked by FDR, Figure 4F), consistent with our SFTPC western blot and immunostaining findings; however, additional AEC2 proteins were also over-represented in the top 50 most upregulated proteins in mutant cells, including SFTPB, SFTPA1, SLPI, and PGC, suggesting that both transcripts and proteins encoding multiple surfactants and other AEC2 markers were in the top most upregulated in mutant iAEC2s. Proteomic analysis also validated the less proliferative state of mutant iAEC2s as MKI67 protein was enriched in corrected cells, whereas cell cycle inhibitors CDKN1C and CDKN2C were upregulated in mutant cells (Figure 4F). Analysis of sections of the patient’s explanted lung also revealed a higher *in vivo* pro-SFTPC immunostaining intensity compared to control distal lung (Figure 4G), suggesting accumulation of mutant pro-SFTPC in the patient’s AEC2s.

Having established that mutant iAEC2s display a less proliferative and more mature AEC2 phenotype than corrected cells with marked accumulation of SFTPC protein, we next sought to determine signaling pathways or biological processes that might differ in mutant cells. The autophagy-related protein ATG2B, essential for lysosomal formation was in the top 10 most upregulated proteins (ranked by logFC) in mutant cells (Figure 4F) and gene set analysis of GO, KEGG, and HALLMARK terms using both proteomic, scRNA-Seq, and bulk RNA-Seq analyses all revealed significant upregulation of lysosomal and autophagy-related terms in mutant cells (Figure 4H and Supplemental Figure 3E). A systems based approach integrating transcriptomic, proteomic, and phosphoproteomic analyses revealed a high degree of concordant regulation with protein secretion processes being perturbed across all datasets consistent with the build up of secretory proteins evident on Western blots and immunostains. In addition, altered apical junctional processes, glycolysis, STAT3 signaling, MTOR signaling, and inflammatory signaling via NF-κB implied additional broad perturbations were present across a wide variety of epithelial, metabolic, and inflammatory cellular processes in mutant iAEC2s, whereas proliferation pathways (E2F and MYC targets) were downregulated (Figure 4I). Together, these data demonstrate that expression of mutant SFTPC drives a transcriptomic program which markedly alters the iAEC2 phenotype and includes alterations in protein expression, quality control, and intracellular signaling.

### SFTPC^I73T^ iAEC2s display upregulation of autophagy with a reduction in autophagic flux

Given the potential effect of the *SFTPC^I73T^* mutation on autophagy suggested by our bioinformatic analyses (Figure 4H, I), we employed a variety of independent static and dynamic approaches to evaluate the autophagy pathway in iAEC2s (Figure 5A). We first assessed the intracellular levels of microtubule-associated protein 1 light chain 3 (LC3), a marker of autophagosome formation (Klionsky et al., 2016). Western blot analysis showed elevated LC3, suggesting an increased autophagosomal mass in the mutant compared to the corrected iAEC2s (Figure 5B). To distinguish between enhanced autophagosome formation vs. reduced degradation, we examined the levels of p62 (SQSTM1). P62 through direct interaction with LC3, facilitates transfer of polyubiquitinated proteins to the completed autophagosome where it is degraded along with its cargo, thus serving as a marker of autophagic degradation. The mutant iAEC2s demonstrated significant accumulation of p62,compared to their corrected counterparts, consistent with a reduction in autophagosome degradation (Figure 5B). To further delineate if the accumulation of LC3 in the mutant iAEC2s was the result of only reduced autophagosome degradation or a concomitant induction in autophagosome formation, we performed autophagic flux studies. We first examined the response to the mTOR inhibitor/autophagy activator torin. When compared to their corrected counterparts, the mutant iAEC2s demonstrated significantly lower LC3 turnover following treatment with torin for 18h (55.2 ± 9.4% vs 26.7 ± 0.3% reduction over baseline, p= 0.05) (Figure 5C). We then examined the effect of bafilomycin A1 (BafA1), a vacuolar-type H+-ATPase (V-ATPase) inhibitor, which has been shown to inhibit autophagosome degradation via both neutralizing the lysosomal pH as well as by blocking the fusion of autophagosomes with lysosomes (Klionsky et al., 2016; Yamamoto et al., 1998). Treatment with BafA1 50nM resulted in earlier accumulation and higher amounts of LC3 in the mutant compared to the corrected iAEC2s suggesting increased autophagosome formation (Figure 5D). The autophagy perturbations observed in the mutant iAEC2s were further validated in the patient line biallelically expressing both the mutant and corrected *SFTPC* alleles (*SFTPC^I73T/WT^*), as evidenced by the accumulation of both LC3 and p62 in the mutant iAEC2s (Supplemental Figure 1B). Together, these results suggest a concomitant upregulation of autophagosome formation (flux) in the mutant iAEC2s as well as a late block in autophagy (autophagosomes turnover) consistent with our previous studies in heterologous cell lines stably expressing SFTPC^I73T^ and *in vivo* in our *SFTPC*^I73T^ mouse model (A. Hawkins et al., 2015; Nureki et al., 2018).

**Figure 5.**
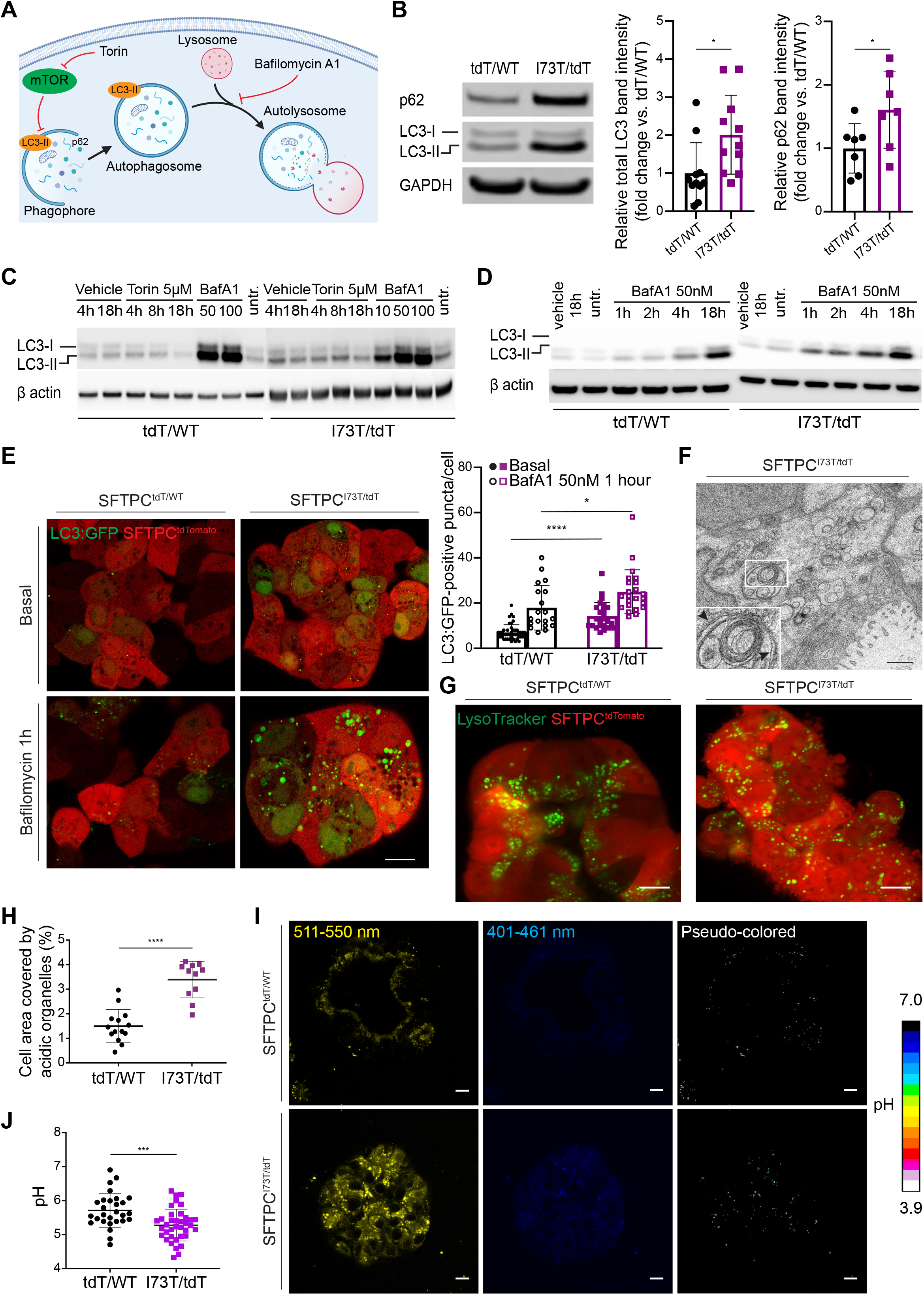
Mutant iAEC2s display upregulation of autophagy but inhibited autophagic flux. (A) Schematic illustrating the autophagy pathway. Key proteins involved in the pathway, such as p62/SQSTM1 and LC3, as well as the mechanisms of action of bafilomycin and torin are depicted. (B) Representative Western blot of SFTPC^tdT/WT^ and SFTPC^I73T/tdT^ iAEC2 lysates for p62 and LC3 with GAPDH and β actin as loading controls. Densitometric quantification showing increased accumulation of both LC3 and p62 in SFTPC^I73T/tdT^ iAEC2s (mean ± SD; n=11 for LC3 and n=7 for p62 independent experiments). (C) Western blots of cell lysates for LC3 and β actin from autophagic flux studies using SFTPC^tdT/WT^ or SFTPC^I73T/tdT^ iAEC2s treated with either torin 5 μM for 4, 8, or 18 hours or bafilomycin A1 (BafA1) 50 nM or 100 nM for 18 hours or vehicle (DMSO) for 4 or 18 hours (blots shown are representative of n=2 independent experiments). (D) A repeat autophagic flux study, as in C, but using bafilomycin A1 (BafA1) 50 nM for 1, 2, 4, and 18 hours or vehicle (DMSO) for 18 hours. SFTPC^I73T/tdT^ iAEC2s accumulate higher amounts of LC3 compared to SFTPC^tdT/WT^ iAEC2s suggesting increased autophagosome formation (representative of n=2 independent experiments). (E) SFTPC^tdT/WT^ and SFTPC^I73T/tdT^ iAEC2s transduced with a lentiviral LC3:GFP fusion protein and exposed to bafilomycin to quantify autophagosomes. Scale bars: 10 μm. (representative confocal fluorescence microscopy images of n=3 biological replicates of independent wells of a differentiation). (F) Representative TEM image of SFTPC^I73T/tdT^ iAEC2s shows a double-membrane autophagosome (inset, arrowheads). Scale bar: 1 μm. (G) Representative live-cell confocal fluorescence microscopy of SFTPC^tdT/WT^ and SFTPC^I73T/tdT^ iAEC2s stained with LysoTracker green, a basic dye that stains acidic vesicular organelles. Scale bars: 10 μm. (H) Quantification of acidic organelles expressed as percentage of covered cell area reveals that SFTPC^I73T/tdT^ iAEC2s contain a higher number of acidic organelles compared to SFTPC^tdT/WT^ iAEC2s. (mean ± SD; n=11-14 independent alveolospheres treated as biological replicates). (I) Representative live-cell confocal fluorescence microscopy of SFTPC^tdT/WT^ and SFTPC^I73T/tdT^ iAEC2s stained with Lysosensor yellow/blue-dextran and imaged using two filter ranges, 401-461 nm (blue) and 511-550 nm (yellow). Subcellular organelles with lower pH are identified as pixels with increased yellow/blue ratio. Pseudo-colored images reflect the organelle pH. Scale bars: 10 μm. (J) Quantification of lysosomal pH reveals that SFTPC^I73T/tdT^ iAEC2 lysosomes demonstrate a lower pH when compared to SFTPC^tdT/WT^ iAEC2 lysosomes. (mean ± SD; n=2 independent experiments with 20-30 cells analyzed per experiment). *p<0.05, ***p<0.001, ****p<0.0001, by unpaired, two-tailed Student’s t-test for all panels.

To further corroborate the observed changes in biochemical markers of autophagy, we next utilized advanced imaging techniques to assess autophagosome dynamics in individual iAEC2s. We first transduced the mutant and corrected iAEC2s with a lentiviral construct constitutively expressing GFP fused to LC3 (LC3:GFP) (Twig et al., 2008) in order to visualize and quantitate autophagosome numbers, which are represented by cytosolic GFP positive punctae. Consistent with the biochemical profile (Figure 5B), under steady state conditions, we found more autophagosomes were present in mutant vs. corrected iAEC2s (Figure 5E). The fusion between autophagosomes and lysosomes promotes the degradation of both LC3 and GFP facilitated by the low pH of the autophagolysosome which can be blocked by treatment with BafA1, a v-ATPase inhibitor which inhibits vacuolar acidification (Ni et al., 2011). Treatment with BafA1 50 nM for 1h enhanced both GFP fluorescence and the numbers of GFP punctae in mutant iAEC2s compared to similarly treated corrected iAEC2 controls consistent with increased autophagic flux (Figure 5E). The changes in autophagosome dynamics in mutant iAEC2s were further confirmed using ultrastructural analysis by TEM where double membrane intracellular vacuoles characteristic of autophagosomes were easily visualized in the mutant iAEC2s, whereas these structures remained sparse in any of the corrected iAEC2s (Figure 5F).

As fusion with acidic lysosomes is required for autophagosome degradation, we considered the possibility that failure of lysosomes to properly acidify might explain the block in autophagy observed in mutant iAEC2s. We therefore stained the mutant and corrected iAEC2s with LysoTracker green, a basic dye that stains acidic vesicular organelles (Figure 5G) and found that the mutant iAEC2s contained more acidic organelles compared to the corrected cells (Figure 5H). To better assess the pH of lysosomes in AEC2s, we next used Lysosensor yellow/ blue-dextran (Figure 5I), a pH-sensitive probe that emits predominantly yellow fluorescence in acidic organelles and blue fluorescence in less acidic organelles and which has been shown to selectively target the lysosomes (Wolfe et al., 2013). Quantitation of probe fluorescence revealed that lysosomes of mutant iAEC2s maintained a lower intraluminal pH when compared to those of their corrected counterparts. Together, these data indicate that the mutant iAEC2s exhibit an accelerated autophagy flux (autophagosome and autophagolysosome formation), without impairment of vacuolar acidification, suggesting that the enhanced production of mutant SFTPCI73T has saturated the autophagic capacity of the mutant iAEC2s without altering lysosomal pH.

### SFTPC^I73T^ iAEC2s demonstrate metabolic reprogramming from oxidative phosphorylation to glycolysis

The recognition that changes in metabolic pathways (i.e. glycolysis) were among the top differentially upregulated pathways in the mutant iAEC2s in our integrated systems analysis (Figure 4H) coupled with our prior work showing that expression of SFTPC^I73T^ could also alter steady state mitochondrial mass and membrane potential (A. Hawkins et al., 2015; Nureki et al., 2018), directed us to next investigate changes in the cellular bioenergetics of mutant iAEC2s. Mitochondrial morphology and mass were assessed using quantitative imaging of MitoTracker dye staining. Distinct mitochondrial morphologies were observed in the mutant vs. corrected iAEC2s. While the mitochondria in the corrected cells were more elongated forming a network extending from the basolateral to the apical cell surface, in the mutant cells the mitochondria demonstrated a rounded, fragmented morphology (Figure 6A). The mutant iAEC2s also demonstrated higher mitochondrial mass (Figure 6B) with accumulation of larger (Figure 6C) but more fragmented mitochondria as assessed by the ratio of the long over the short diameter (aspect ratio) (Figure 6D), changes previously described in primary AEC2s from IPF patients and aged mice (Bueno et al., 2015). The higher mitochondrial mass in the mutant iAEC2s was confirmed by higher levels of the mitochondrial protein TOM20, which also appeared to be time-dependent with enhanced accumulation as the cells aged in culture (Figure 6E).

**Figure 6.**
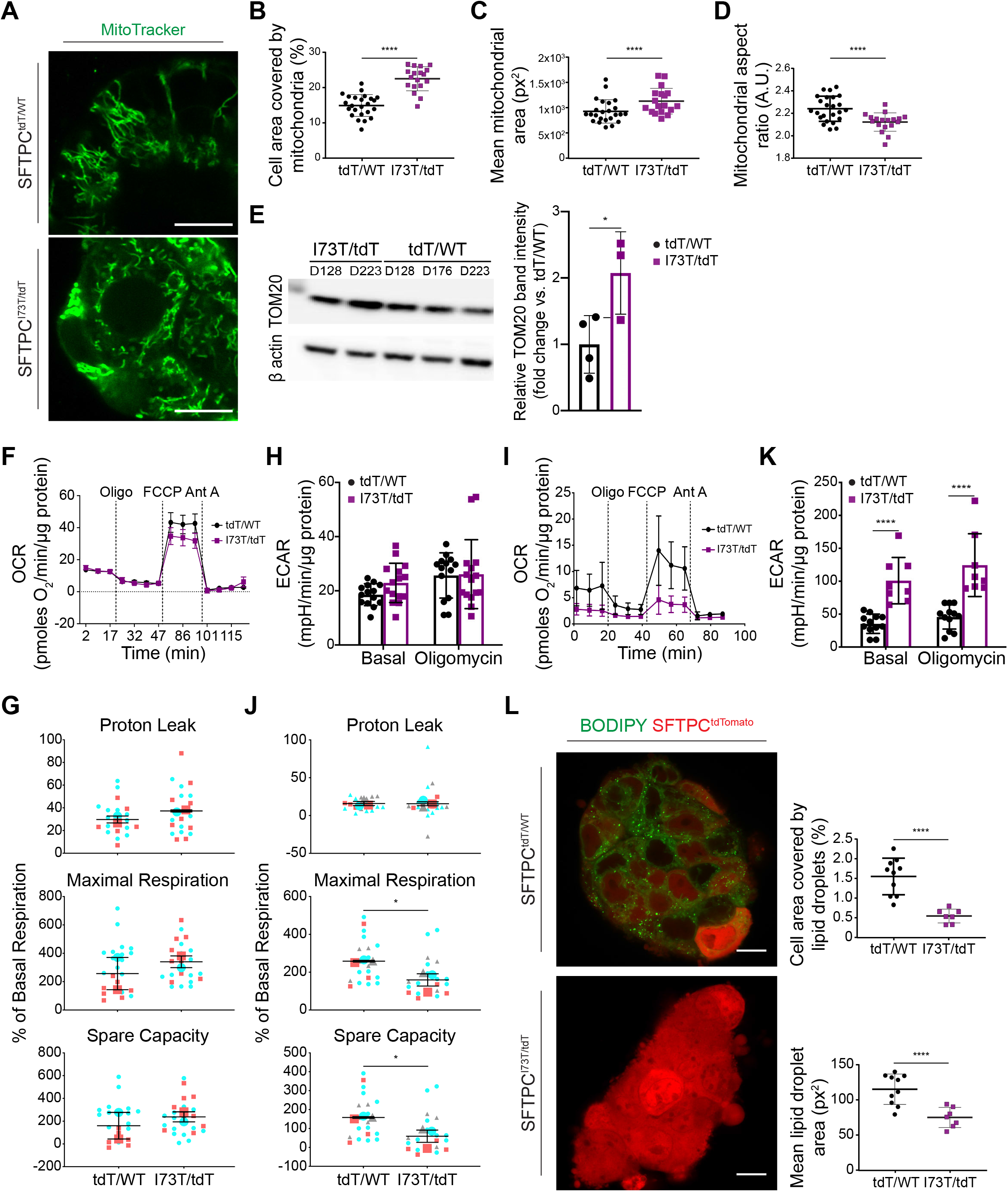
SFTPCI73T expression leads to accumulation of dysfunctional mitochondria and metabolic reprogramming from oxidative phosphorylation to glycolysis. (A) Representative live-cell confocal fluorescence microscopy of SFTPC^tdT/WT^ and SFTPC^I73T/tdT^ iAEC2s stained with MitoTracker green shows distinct mitochondrial morphologies. Scale bars: 10 μm. (B-D) Quantitative analyses of morphometric data from fluorescence images. (B) Increased mitochondrial mass in SFTPC^I73T/tdT^ iAEC2s, quantified as the percentage of cell area covered by mitochondria. (C) Increased mitochondrial size in SFTPC^I73T/tdT^ iAEC2s. (D) SFTPC^I73T/tdT^ iAEC2 mitochondria are more fragmented, assessed by the aspect ratio. (B-D data represent mean ± SD; n=24 SFTPC^tdT/WT^ and n=18 SFTPC^I73T/tdT^ independent alveolospheres treated as biological replicates). (E) Representative Western blot of SFTPCtdT/WT and SFTPCI73T/tdT iAEC2 lysates for TOM20 with β actin as loading control. Densitometric quantification showing increased mitochondrial mass in SFTPCI73T/tdT iAEC2s (mean ± SD; n=4 (days 128, 176, 223, 223) SFTPCtdT/WT and n=3 (days 128, 223, 223) SFTPCI73T/tdT independent experiments). (F) Early time point OCR was measured under basal conditions followed by addition of oligomycin (4.5 μM/l), FCCP (1 μM/l), and antimycin A (Ant A; 2.5 μM/l), as indicated. (G) SFTPC^tdT/WT^ and SFTPC^I73T/tdT^ iAEC2s demonstrate no significant differences in proton leak, maximal respiration, and spare capacity measured as percent change over basal respiration in the early time point (superplots where small shapes represent replicate values within each experiment and large shapes represent the average of each independent experiment, tdT/WT and I73T/tdT are colored-matched between experiments, mean ± SEM; n=2 independent experiments). (H) Early time point ECAR was measured under basal conditions followed by addition of oligomycin (4.5 μM/l), as indicated, shows no significant differences between SFTPCtdT/WT and SFTPCI73T/tdT iAEC2s. (I) Late time point OCR was measured under basal conditions followed by addition of oligomycin (4.5 μM/l), FCCP (1 μM/l), and antimycin A (Ant A; 2.5 μM/l), as indicated. (J) SFTPCI73T/tdT iAEC2s demonstrate significantly lower maximal respiration and spare respiratory capacity (superplots where small shapes represent replicate values within each experiment and large shapes represent the average of each independent experiment, tdT/WT and I73T/tdT are colored-matched between experiments, mean ± SEM; n=3 independent experiments) measured as percent change over basal respiration. (K) Late time point ECAR was measured under basal conditions followed by addition of oligomycin (4.5 μM/l), as indicated, revealing a significantly higher ECAR in SFTPCI73T/tdT iAEC2s. (mean ± SD; n=3 independent experiments). (L) Representative live-cell confocal fluorescence microscopy of SFTPCtdT/WT and SFTPCI73T/tdT iAEC2s stained with BODIPY 493/503 dye. Quantitative analyses reveal a significantly lower amount and smaller size of neutral lipid droplets in SFTPCI73T/tdT iAEC2s. Scale bars: 10 μm. (mean ± SD; n=10 SFTPCtdT/WT and n=7 SFTPCI73T/tdT independent alveolospheres treated as biological replicates). *p<0.05, ****p<0.0001 by unpaired, two-tailed Student’s t-test for all panels.

To investigate whether the changes in mitochondrial morphology between the mutant and corrected iAEC2s were associated with changes in mitochondrial function, we performed respirometry experiments at different time points: earlier time points close to our prior transcriptomic/proteomic analyses (differentiation days 128 and 153) and later time points (differentiation days 175, 213, and 291). At the earlier time points, there were no significant differences between the mutant and corrected iAEC2s in terms of their basal oxygen consumption rate (OCR), maximal uncoupled mitochondrial respiration (assessed after the administration of FCCP), and spare respiratory capacity (Figure 6F, G). Similar to oxidative phosphorylation, no differences were observed in glycolysis between the mutant and corrected iAEC2s, as measured by the extracellular acidification rate (ECAR) either at steady-state or following treatment with the ATP synthase inhibitor oligomycin (Figure 6H). However, at later time-points, which correspond to further aged iAEC2s, the mutant cells demonstrated a significant reduction in maximal uncoupled mitochondrial respiration and spare respiratory capacity (Figure 6I, J). Conversely, the mutant iAEC2s at later time points demonstrated a higher ECAR (Figure 6K) suggestive of a metabolic switch towards glycolytic metabolism. As mobilization of lipid stores is required not only for energy utilization but also to sustain the expansion of the surface area of the mitochondrial membranes and autophagosomes, we assessed the lipid content of mutant and corrected iAEC2s. We found significantly lower levels of stored triglycerides in the mutant iAEC2s, as assessed by BODIPY493/503 dye (Figure 6L). Together, these data suggest that *SFTPC* mutant iAEC2s demonstrate metabolic reprogramming from oxidative phosphorylation to anaerobic metabolism and have depleted lipid stores, a process that appears to be time-dependent.

### SFTPCI73T iAEC2s elicit an inflammatory response

The proteomic analysis revealed upregulation of the NF-κB pathway in the SFTPCI73T mutant iAEC2s compared to their corrected counterparts (Figure 7A). Since autophagy inhibition has been shown to lead to activation of the NF-κB pathway via upregulation of p62 (Hill et al., 2019), we sought to further validate activation of the NF-κB pathway in mutant iAEC2s imputed by the proteomic analysisTo do this we utilized a lentiviral vector (lenti-NF-κB-luc-GFP) (Figure 7B) that we previously engineered to enable independent simultaneous tracking of transduced (GFP+) cells by flow cytometry and real-time assessment of NF-κB activation levels quantified by a luciferase reporter whose expression is regulated by a minimal promoter element preceded by consensus binding sites for the canonical NF-κB p50-p65 heterodimer transcription factor complex (Wilson et al., 2013).Mutant and corrected iAEC2s were infected with the lentiviral vector (MOI 20: Figure 7C), allowed to expand in culture, and transduced (GFP+) SFTPC^tdTomato+^ mutant and corrected iAEC2s were then sorted and analyzed for luciferase expression (Figure 7D). The *SFTPC^I73T^* mutant iAEC2s demonstrated increased bioluminescence when compared to their corrected counterparts, suggesting increased canonical NF-κB activity in the mutant iAEC2s (Figure 7E). Furthermore, mutant iAEC2s secreted higher levels of the NF-κB target proteins GM-CSF, CXCL5, and MMP-1 as assessed by Luminex analysis of supernatants, when compared to their corrected counterparts (Figure 7F). Taken together, these findings suggest mutant iAEC2s are an important proinflammatory hub with resultant secretion of inflammatory mediators via activation of the NF-κB pathway.

**Figure 7.**
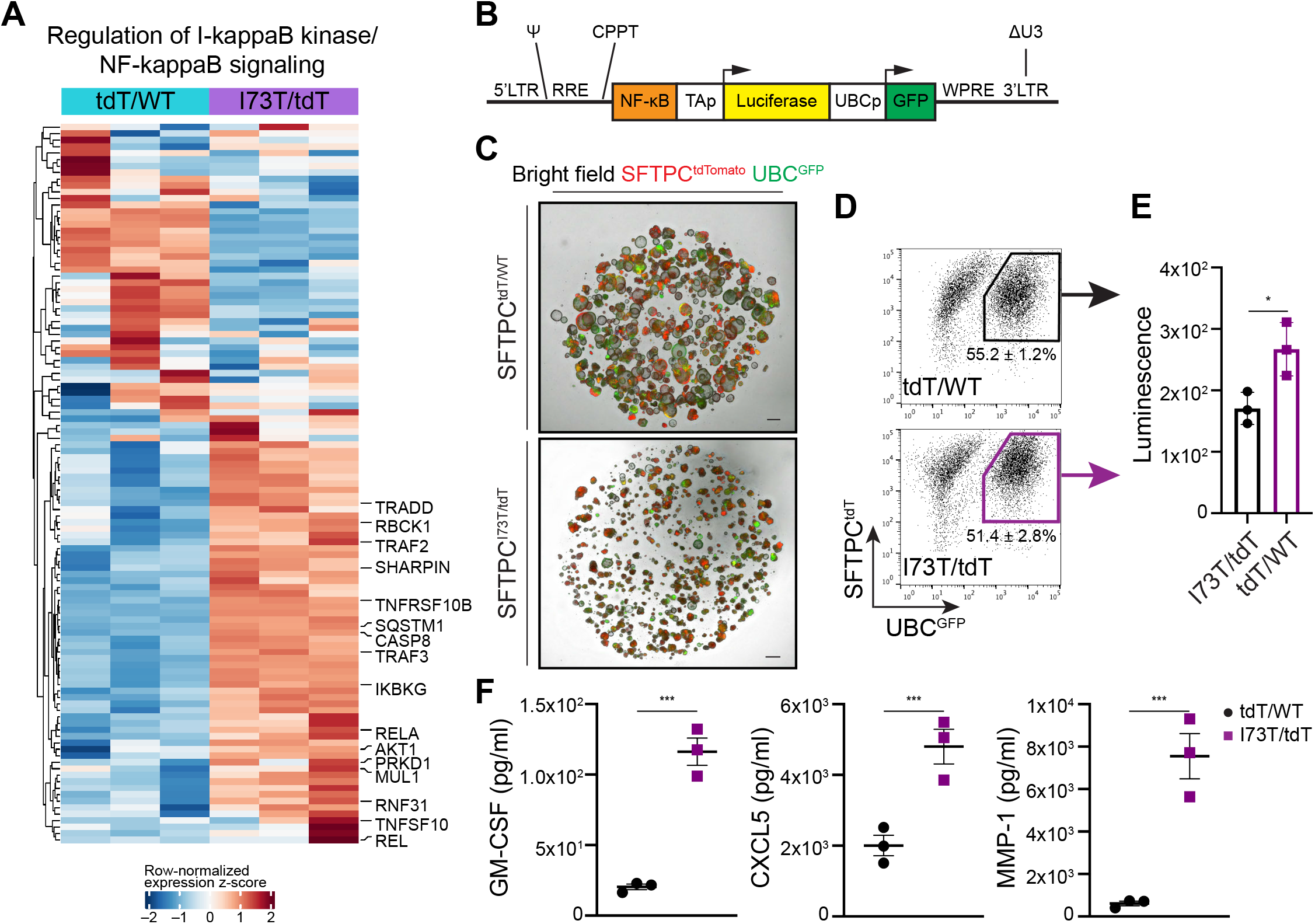
Mutant iAEC2s display activation of the NF-κB pathway. (A) Unsupervised hierarchical clustered heat map of differentially expressed proteins (FDR<0.05) in the GO Biological Process gene set “regulation of I-kappaB kinase/NF-kappaB signaling”, as plotted with row normalized z-score; a selected subset of differentially expressed proteins is highlighted with large font. (B) Schematic of NF-κB-luc-GFP lentiviral construct: four copies of the canonical NF-κB p50/p65 heterodimer consensus binding sequence precede the minimal thymidine kinase promoter (TAp) of the herpes simplex virus. Promoter activation in the presence of NF-κB signaling drives expression of the firefly luciferase reporter. GFP is constitutively expressed by the ubiquitin C (UBC) promoter, allowing purification of transduced cells by FACS. LTR: lentiviral long terminal repeats, RRE: rev responsive element, CPPT: central polypurine tract, WPRE: Woodchuck hepatitis virus post-transcriptional regulatory element, ΔU3: deleted U3 region for in vivo inactivation of the viral LTR promoter, Ψ: Psi lentiviral packaging sequence, GFP: green fluorescence protein. (C) Representative live-cell imaging of lentivirally transduced SFTPC^tdT/WT^ and SFTPC^I73T/tdT^ alveolospheres (bright-field/tdTomato/ GFP overlay). Scale bars: 500 μm. (D) Sort gates used to purify transduced (GFP+) SFTPC^tdT/WT^ and SFTPC^I73T/tdT^ iAEC2s. (E) Bioluminescence quantification shows increased luciferase activity in SFTPC^I73T/tdT^ compared to SFTPC^tdT/WT^ iAEC2s. (mean ± SD; n=3 biological replicates of independent wells of a differentiation). (F) Luminex analysis of supernatants collected from SFTPC^I73T/tdT^ and SFTPC^tdT/WT^ iAEC2 cultures reveals higher concentrations of the NF-κB target proteins GM-CSF, CXCL5, and MMP-1 in the SFTPC^I73T/tdT^ iAEC2 cultures. (mean ± SD; n=3 biological replicates of independent wells of a differentiation). *p<0.05, **p<0.01, ***p<0.001 by unpaired, two-tailed Student’s t-test for all panels.

### Treatment of SFTPC^I73T^ iAEC2s with hydroxychloroquine results in aggravation of autophagy perturbations and metabolic reprogramming

To apply our iPSC-derived model system as a preclinical disease model for drug testing, in a proof-of-principle experiment we assessed the effect of hydroxychloroquine (HCQ) on *SFTPC^I73T^* expressing iAEC2s. HCQ is a medication commonly used either alone or in combination with corticosteroids in pediatric patients with genetic disorders of surfactant dysfunction affecting AEC2s (including SFTPC^I73T^) and resulting in chILD. The use of HCQ in this patient population is based primarily on evidence from case reports or small cohort studies with variable results (Braun et al., 2014; Klay et al., 2018; Kröner et al., 2015). HCQ is a lysosomotropic medication that accumulates in lysosomes inhibiting lysosomal activity and autophagy (Mauthe et al., 2018; Schrezenmeier and Dörner, 2020) raising the possibility it may aggravate the autophagic block already present in *SFTPC^I73T^* expressing iAEC2s. Indeed, treatment of *SFTPC^I73T^* expressing iAEC2s with HCQ (10 μM) for 7 days resulted in increased accumulation of LC3 in HCQ-treated compared to untreated iAEC2s (Figure 8A). Notably, treatment of the corrected iAEC2s with HCQ at the same concentration and for the same duration did not result in significant changes of total LC3 suggesting that mutant iAEC2s are more sensitive to HCQ’s lysosomotropic effect. Long-term treatment of *SFTPC^I73T^* expressing iAEC2s with HCQ (10 μM) resulted in further reduction of the already diminished self-renewal capacity of these cells (Figure 8B), while both mutant and corrected iAEC2s retained expression of the SFTPC^tdTomato^ reporter (Figure 8C).

**Figure 8.**
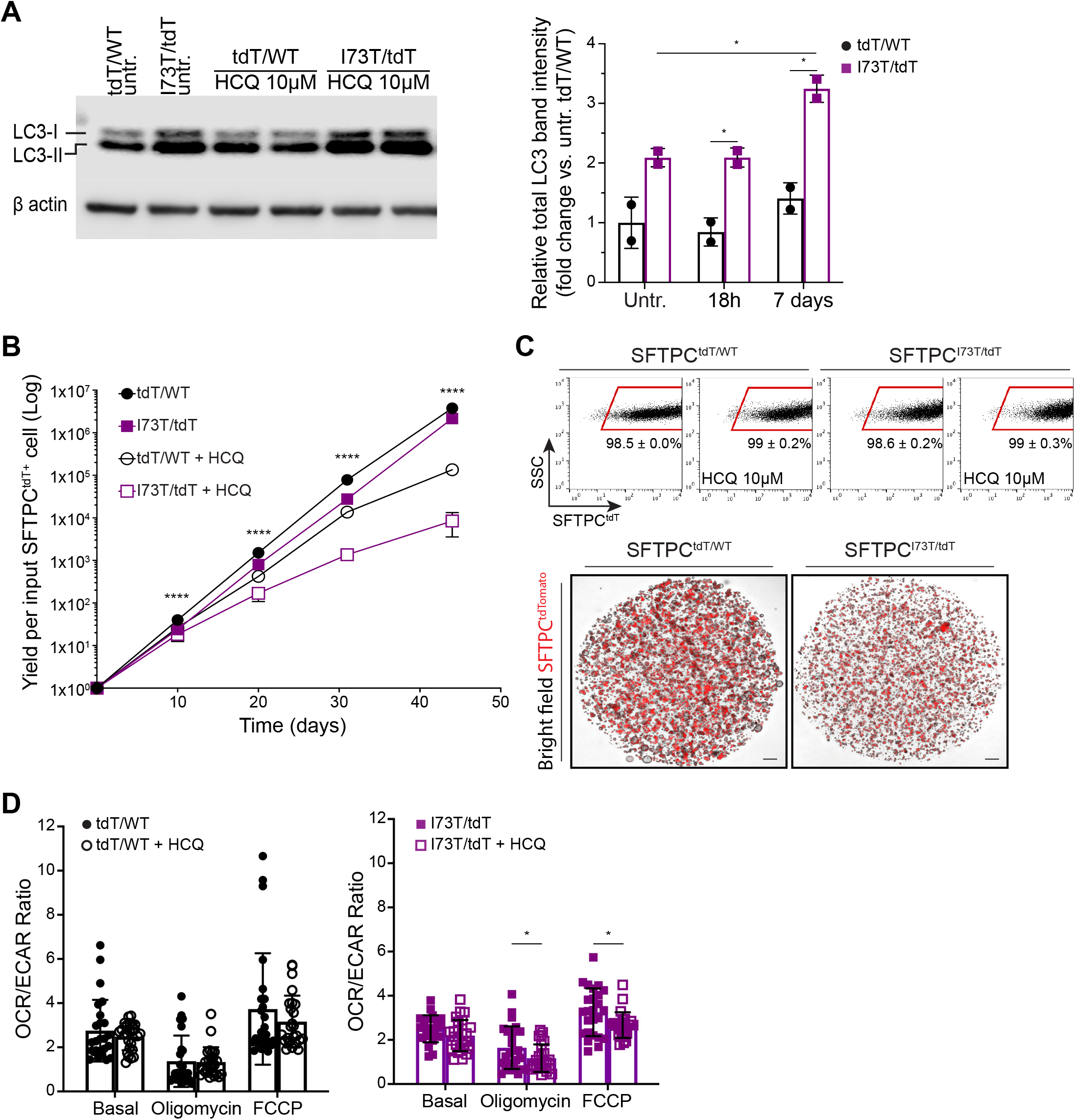
Hydroxychloroquine treatment aggravates the autophagic flux block and metabolic reprogramming in *SFTPC^I73T^* expressing iAEC2s. (A) SFTPC^tdT/WT^ or SFTPC^I73T/tdT^ iAEC2s were treated with either hydroxychloroquine (HCQ; 10 μM) or vehicle (ddH2O) for 18 hours or 7 days. Lysates were subjected to Western blotting for LC3 with β actin as loading control. A representative Western blot following HCQ treatment for 7 days is shown (n=2 biological replicates of independent wells of a differentiation). Densitometric quantification showing increased accumulation of LC3 in SFTPC^I73T/tdT^ iAEC2s treated with HCQ for 18 hours or 7 days compared to untreated SFTPC^I73T/tdT^ iAEC2s as well as compared to their corrected SFTPC^tdT/ WT^ counterparts treated with HCQ for the same duration. (n=2 biological replicates of independent wells of a differentiation). (B) Graph showing yield in cell number per input SFTPC^tdT/WT^ or SFTPC^I73T/tdT^ iAEC2 with or without treatment with HCQ (10 μM). (mean ± SD; n=3 biological replicates of independent wells of a differentiation). ****p<0.0001 by one-way ANOVA. (C) Representative flow cytometry dot plots of SFTPC^tdT/WT^ and SFTPC^I73T/tdT^ iAEC2s with or without HCQ (10 μM) treatment and representative live-cell imaging of SFTPC^tdT/WT^ and SFTPC^I73T/tdT^ HCQ-treated alveolospheres (bright-field/tdTomato overlay) showing maintenance of the AEC2 program (mean ± SD is shown; n=3 biological replicates of independent wells of a differentiation). Scale bars: 500 μm. (D) Representative bioenergetics bar graphs plotted as OCR/ECAR ratios under basal conditions or following addition of oligomycin (4.5 μM/l) or FCCP (1 μM/l) for SFTPC^tdT/WT^ (black circles) and SFTPC^I73T/tdT^ (purple squares) iAEC2s with or without HCQ (10 μM) treatment for 7 days show a significant reduction in OCR/ECAR ratio only in SFTPC^I73T/tdT^ iAEC2s treated with HCQ (n=2 independent experiments). *p<0.05, by unpaired, two-tailed Student’s t-test for all panels.

Furthermore, *SFTPC^I73T^* expressing iAEC2s treated with HCQ for 7 days demonstrated a significant reduction in OCR/ECAR ratio assessed after the administration of oligomycin and FCCP when compared to untreated *SFTPC^I73T^* iAEC2s, suggesting a shift toward anaerobic metabolism revealed after pharmacological manipulation of energy supply and demand, and reminiscent of the observations made earlier for aged mutant iAEC2s (Figure 6). Similar to LC3 levels, HCQ treatment had no effect on the OCR/ECAR ratio in the corrected iAEC2s (Figure 8D).

## Discussion

Epithelial cell dysfunction has been implicated as an upstream driver in the pathogenesis of IPF; however, understanding the cellular and molecular mechanisms by which dysfunctional AEC2s initiate the fibrotic cascade in humans has been hindered by difficulty in accessing patient samples at an early disease stage and the inability to maintain primary AEC2s in culture for prolonged periods. Despite the recent progress made by the application of genetic mouse models (Katzen et al., 2019; Nureki et al., 2018) and the interrogation of fibrotic human lungs with contemporary molecular methods (Adams et al., 2020; Habermann et al., 2020; Reyfman et al., 2018), the significant differences between murine and human pulmonary fibrosis, as well as the advanced fibrosis present in the analyzed human samples (which makes the distinction between primary and secondary effects difficult) highlight the need for reliable human preclinical disease models to study the intrinsic epithelial cell dysfunction at the inception of the fibrotic process.

Our results herein suggest that patient-specific iAEC2s can serve as a human preclinical disease model system, not only unraveling the pathophysiology of human AEC2 dysfunction but also providing a novel platform to test currently used unproven or novel therapies. Patient-derived iAEC2s expressing mutant SFTPC^I73T^ demonstrated misprocessing and mistrafficking of pro-SFTPC protein to the plasma membrane recapitulating the mistrafficking occurring in the donor patient’s in vivo AEC2s. While the accumulation of misprocessed pro-SFTPC forms, as a result of the *SFTPC^I73T^* mutation, has been previously shown in overexpression experiments in heterologous cell lines (Brasch et al., 2004; Galetskiy et al., 2008; A. Hawkins et al., 2015; Stewart et al., 2012), patient bronchoalveolar lavage samples (Brasch et al., 2004), and most recently in an in vivo mouse model (Nureki et al., 2018), to our knowledge this represents the first human model in which *SFTPC^I73T^* is expressed from the endogenous locus.

The iAEC2 platform also offers the ability to temporally model both the early changes coinciding with accumulation of mutant SFTPC forms and the resulting downstream sequence of events that result in the acquisition of a dysfunctional AEC2 phenotype. Multi-omic profiling of the mutant and corrected iAEC2s identified similarities at an early time point but more distinct phenotypes emerged with culture expansion and serial passaging suggesting a time-dependent phenotype. This time course is in agreement with the hypomorphic *SFTPC^I73T^* mouse, where an age-dependent lung phenotype emerged simultaneously with the gradual temporal accumulation of mutant pro-SFTPC forms (Nureki et al., 2018). Although both corrected and mutant iAEC2s had similar frequency of expression of the AEC2 program, mutant iAEC2s were less proliferative and demonstrated higher expression of AEC2 marker genes especially at later time points.

Among the many cellular programs disrupted in the iAEC2 model, our findings suggest that accumulation of misprocessed and mistrafficked pro-SFTPC results in an early perturbation in autophagy. First, unbiased signaling pathway analysis identified the lysosomal/autophagy pathway as being differentially upregulated in the mutant iAEC2s. We next validated these observations by a variety of independent static and dynamic approaches (Figure 5) which documented induction of autophagosome formation (flux) but defined a late block in autophagy at the level of autophagosomes turnover, as evidenced by the accumulation of both LC3 and p62 and the incomplete response to torin in the mutant iAEC2s. Our results are consistent with previous studies in heterologous cell lines stably expressing SFTPC^I73T^ and in vivo in our *Sftpc^I73T^* mouse model (A. Hawkins et al., 2015; Nureki et al., 2018). Autophagy is an evolutionarily conserved homeostatic process during which cells degrade and recycle unnecessary or dysfunctional organelles and proteins via the formation of double-membraned autophagosomes which fuse with lysosomes (Klionsky et al., 2016). A dysregulated autophagic response has been previously shown in IPF lungs, evidenced by accumulation of p62 (SQSTM1) (Araya et al., 2013; Hill et al., 2019; Patel et al., 2012), and has been postulated to subsequently lead to epithelial cell senescence and myofibroblast differentiation (Araya et al., 2013; Hill et al., 2019). Although it remains unclear mechanistically how SFTPC buildup/ misprocessing leads to autophagy perturbations, we speculate that it is likely related to failure of lysosomal fusion as lysosomal acidification appears intact in *SFTPC^I73T^* expressing iAEC2s. The resultant proteostatic perturbations and alterations in cellular quality control resulted in diminished progenitor potential as evidenced by reduced self-renewal capacity. Selective ablation of AEC2s has been previously shown to promote either spontaneous lung fibrosis (O. Garcia et al., 2016; Sisson et al., 2010) or enhanced susceptibility to intra-tracheally administered bleomycin (Barkauskas et al., 2013). More recently, senescence induction via conditional deletion of Sin3a in adult mouse AEC2s resulted in progressive pulmonary fibrosis (Yao et al., 2020). Therefore, our results are in agreement with the growing body of literature suggesting that dysfunctional AEC2s could initiate the fibrotic cascade.

An important observation in our study is the accumulation of dysfunctional mitochondria in mutant iAEC2s and the resulting metabolic reprogramming from oxidative phosphorylation to glycolysis. To meet the high metabolic demands required for their complex function, AEC2s have a higher mitochondrial mass compared to other lung cell types (Massaro et al., 1975). Furthermore, autophagy plays an important role in the selective degradation of dysfunctional mitochondria. It is therefore not surprising that mutant iAEC2s displaying an autophagy phenotype also accumulated dysmorphic and dysfunctional mitochondria in a time-dependent manner, changes previously described in primary AEC2s from IPF patients and aged mice (Bueno et al., 2015). To our knowledge, our study is the first to assess the bioenergetics of human AEC2s which has not been possible before given the inability to culture primary AEC2s.

In addition to detailed alterations in cell quality control and metabolism, our data also provide a plausible explanation of how a dysfunctional AEC2 signals to surrounding cells to initiate inflammatory and profibrotic cascades, as the NF-κB pathway was found to be differentially upregulated in the mutant compared to corrected iAEC2s. Notably, autophagy inhibition has been shown to lead to activation of the NF-κB pathway via p62 upregulation (Hill et al., 2019). Furthermore, higher levels of the NF-κB target proteins GM-CSF, CXCL5, and MMP-1 were identified in the supernatants of mutant compared to corrected iAEC2s. Importantly, GM-CSF is a strong chemoattractant for profibrotic Ly6c^hi^ monocytes which have been recently implicated in the pathogenesis of lung fibrosis (Misharin et al., 2017; Venosa et al., 2019).

Finally, in a proof-of-principle experiment we tested the application of our model system as a preclinical human disease drug testing platform by assessing the effect of HCQ on *SFTPC^I73T^* expressing cells. HCQ has been commonly used in pediatric patients with genetic disorders of surfactant dysfunction affecting AEC2s and resulting in chILD. Treatment of *SFTPC^I73T^*-expressing iAEC2s in our model resulted in aggravation of the observed autophagy perturbations, further reduction of the already diminished self-renewal capacity and metabolic reprogramming towards anaerobic metabolism. Taken together, the findings from our epithelial-only model system suggest that treatment with HCQ could be detrimental to patients carrying the *SFTPC^I73T^* mutation, although a pleiotropic effect resulting from HCQ’s effect on immune cells or beneficial effects on other cell types *in vivo* cannot be excluded.

Since AEC2 dysfunction resulting from SFTPC misprocessing was evident in our model, a therapeutic approach beyond gene editing is a next logical step for developing new therapies. Gene editing to ablate expression of mutant *SFTPC* restored normal AEC2 function in our patient-specific iAEC2s in vitro as well as in recent work demonstrating successful *in vivo* editing in the lungs of fetal *SFTPC^I73T^* knock-in mice (Alapati et al., 2019). However, a variety of drug approaches including the use of nanoparticles to augment lysosomal acidification or forced over-expression of TFEB, a driver of lysosomal biogenesis, failed to correct the iAEC2 dysfunction in our hands (data not shown), consistent with our findings that lysosomal-autophagosomal acidification is likely intact in mutant *SFTPC^I73T^* iAEC2s. More directed therapies to correct the likely autophagosomal-lysosomal fusion and turnover defects in this phenotype will be required as new data defining its mechanistic biology emerges.

In summary, our epithelial-only patient-specific iAEC2 model system not only recapitulates key observations made previously in heterologous cell lines and in vivo mouse models as well as in vivo in the patient from whom the iPSC lines were generated, but also provides new insights into the potential role of AEC2s in the inception of pulmonary fibrosis. It also provides a novel preclinical platform to test currently used unproven as well as novel therapies.

## Author Contributions

KDA, MFB, and DNK conceived the work. KDA, AP, ET, AE, SHG, OSS, MFB, and DNK designed experiments. KDA, SJR, AP, ET, RAP, SM, RMH, MV, and SK conducted experiments and analyzed data. JCJ and DNK designed the targeting strategy and generated the iPSC lines. CVM and BCB performed bioinformatics analysis. JAW, FSC, and AH provided the patient’s skin fibroblasts and explant lung slides. SHG provided human fetal lung samples. KDA, MFB, and DNK prepared the original draft and KDA, DNK, MFB, SHG, AP, EPT, FSC, JAW, BCB, and RMH edited the manuscript. All authors reviewed and approved the final version prior to submission.

## Acknowledgements

The authors wish to thank all members of the Kotton, Beers, Emili, and Shirihai Labs for insightful discussions. We thank Anne Hinds of the Boston University Pulmonary Center for histology technical support. We thank Yuriy Alekseyev of the Boston University Schoolof Medicine (BUSM) Single Cell Sequencing Core, supported by NIH grant 1UL1TR001430, and Brian R. Tilton of the BUSM Flow Cytometry Core. We are grateful to Greg Miller and Marianne James of the Boston University Center for Regenerative Medicine (CReM) for maintenance and characterization of patient-specific iPSCs, supported by NIH grants NO1 75N92020C00005 and U01TR001810. We thank Michael Morley and Edward E. Morrisey of the University of Pennsylvania for access to their bioinformatics portal for analyses of bulk RNA-seq datasets. We thank Jialiu Zeng and Mark W. Grinstaff of the Boston University Biomedical Engineering Department for kindly providing the photoactivatable lysosome-targeted acidifying nanoparticles. The schematic illustrating the autophagy pathway and the mouse embryo and human child cartoons were created with BioRender. This work was supported by the I.M. Rosenzweig Junior Investigator Award from The Pulmonary Fibrosis Foundation to K.D.A.; NIH grants U01HL148692, U01HL134745, U01HL134766 and R01HL095993, and an IDEAL Consortium Grant from Celgene/Bristol Myers Squibb to D.N.K.; the Albert M. Rose Established Investigator Award from the Pulmonary Fibrosis Foundation, Department of Veterans Affairs Merit Review 1I01BX001176 and NIH grants RO1 HL119436 to M.F.B.; AE acknowledges generous startup funding to the operations of the CNSB from Boston University.

---

**Supplemental Figure 1.**
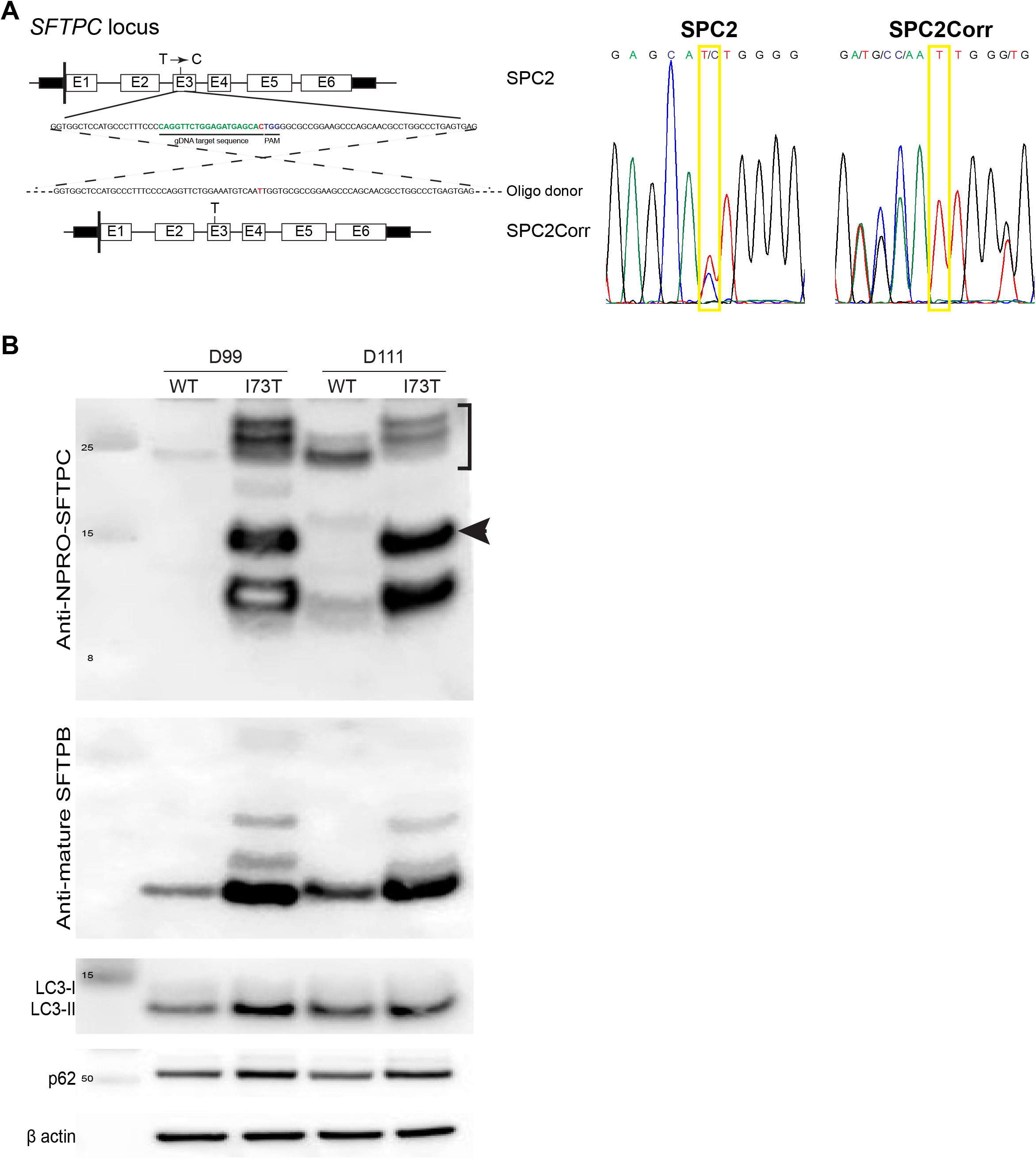
Pro-SFTPC misprocessing and autophagy perturbations in iAEC2s derived from the SFTPC^I73T/WT^ line are ameliorated after gene correction via “footprint-free” CRISPR editing of iPSCs. (A) Footprint free correction of the I73T mutation in *SFTPC* exon 3: genomic sequence with the c.218 T>C (p.I73T) mutation shown in red, the CRISPR guide RNA target sequence shown in green, the protospacer adjacent motif (PAM) cutting site in blue, and the oligo-based donor design with corrected base shown in red. Pre- and post-correction DNA sequencing chromatograms with yellow boxes show the c.218 mutation site sequence. (B) Representative Western blot of SFTPC^WT/WT^ and SFTPC^I73T/WT^ iAEC2 lysates at the indicated time points for pro-SFTPC (NPRO-SFTPC), mature SFTPB, p62 and LC3 with β actin as loading control. SFTPC^I73T/WT^ iAEC2s accumulate large amounts of the primary translation product (bracket) and misprocessed intermediate forms (arrowhead). In SFTPC^WT/WT^ iAEC2s, both the primary translation product and major processing intermediate forms are detected. SFTPC^I73T/WT^ iAEC2s accumulate a larger amount of mature 8 kDa SFTPB. SFTPC^I73T/WT^ iAEC2s demonstrate increased accumulation of both LC3 and p62 compared to SFTPC^WT/WT^ iAEC2s, similarly to SFTPC^I73T/tdT^ iAEC2s (Figure 5B).

**Supplemental Figure 2.**
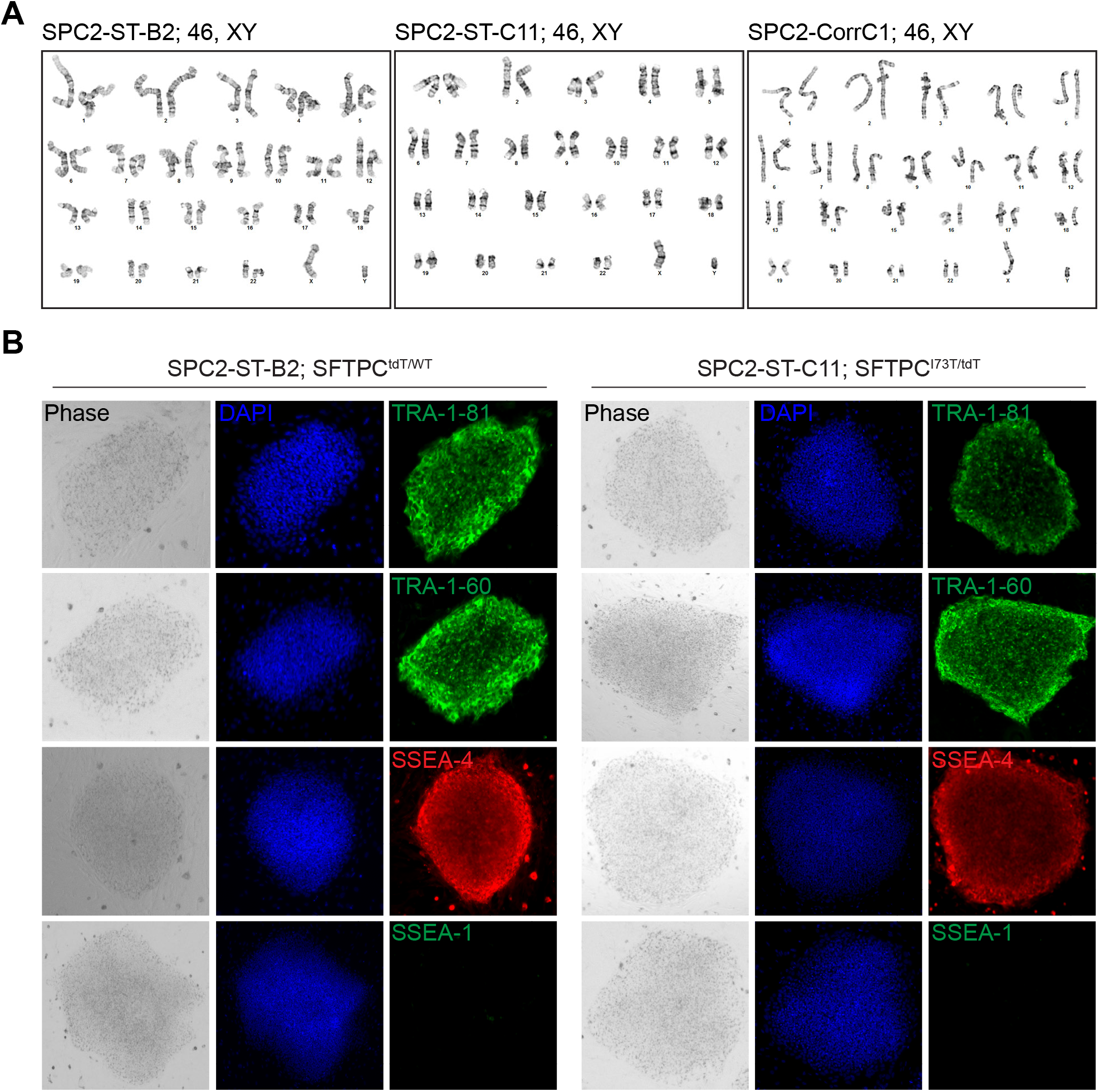
iPSC line characterization. (A) Representative karyotypes (SPC2-ST-C11, SPC2-ST-B2, SPC2-CorrC1 iPSC lines) after gene editing showing a normal 46XY karyotype. (B) Representative immunofluorescence microscopy images showing expression of the pluripotency markers TRA-1-60, TRA-1-81, and SSEA-4 but not SSEA-1, as expected for human pluripotent stem cells (ES Cell Characterization Kit, SCR001 EMD Millipore).

**Supplemental Figure 3.**
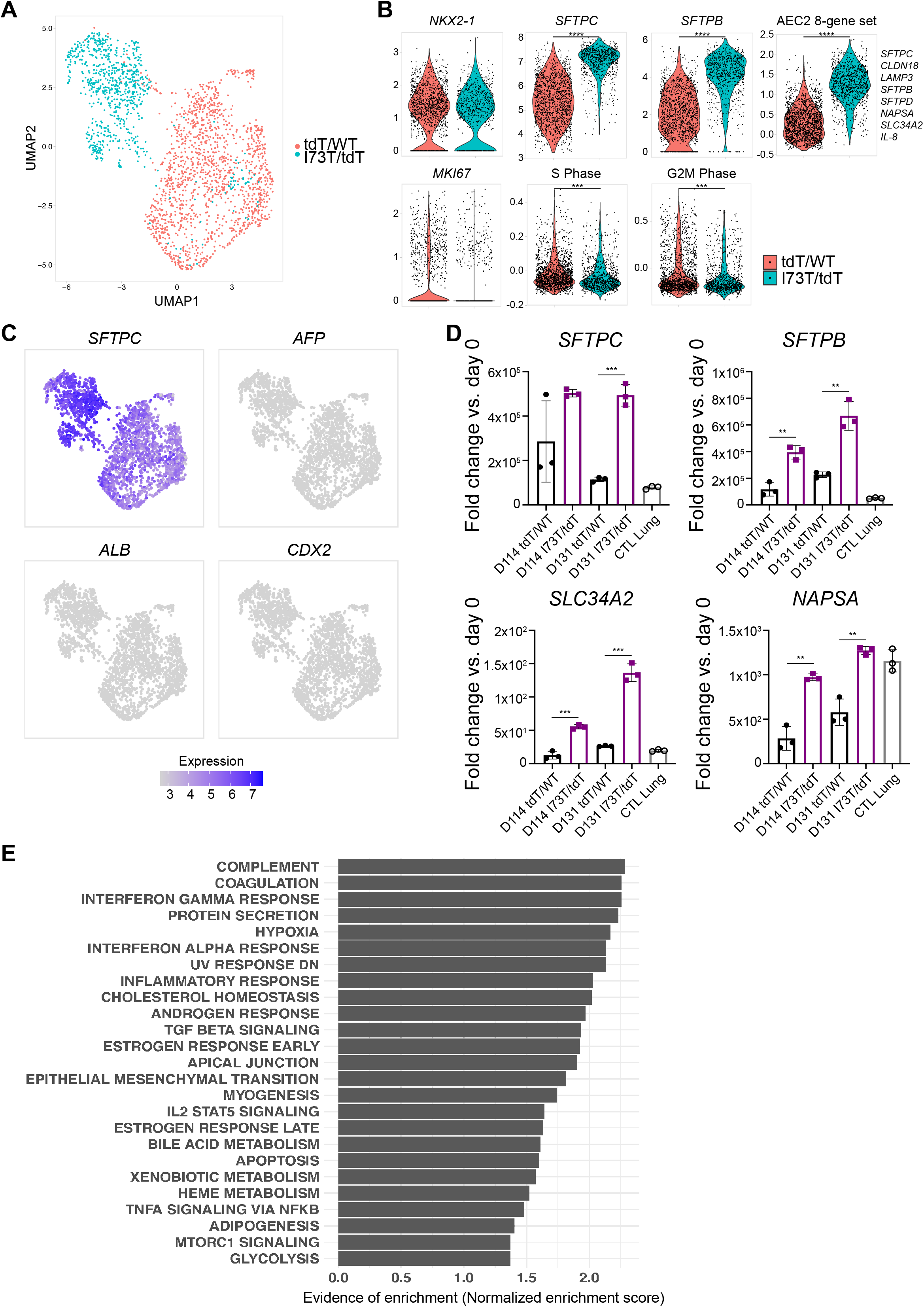
Characterization of gene expression in SFTPC^I73T/tdT^ iAEC2s reveals pathway perturbations and higher expression of AEC2 marker genes. (A) Clustering of day 114 scRNA-seq transcriptomes in each indicated iAEC2 population using Uniform Manifold Approximation Projection (UMAP). (B) Violin plots showing normalized expression for indicated genes or cell cycle phase in day 114 SFTPC^tdT/WT^ and SFTPC^I73T/tdT^ iAEC2s by scRNA-seq. (C) Normalized gene expression overlayed on UMAP plots for SFTPC or each indicated non-lung endodermal transcript. (D) qRT-PCR showing fold change in gene expression compared to day 0 (2^(-ΔΔCt)^) in day 114 and day 131 SFTPC^tdT/WT^ and SFTPC^I73T/tdT^ iAEC2s as well as control samples of an adult control human distal lung explant (CTL Lung). (mean ± SD; n=3 biological replicates of independent wells of a differentiation). (E) Gene set enrichment analysis (GSEA, Camera of day 113 scRNA-seq from Figure 4A, using HALLMARK gene sets) of significantly upregulated gene sets (FDR<0.05) in SFTPC^I73T/tdT^ compared to SFTPC^tdT/WT^ iAEC2s. **p<0.01, ***p<0.001, ****p<0.0001 by unpaired, two-tailed Student’s t-test for all panels.

## Notes

**Conflict of interest:** The authors have declared that no conflict of interest exists.

### Competing Interest Statement

The authors have declared no competing interest.

